# TrancriptomeReconstructoR: data-driven annotation of complex transcriptomes

**DOI:** 10.1101/2020.12.10.418897

**Authors:** Maxim Ivanov, Albin Sandelin, Sebastian Marquardt

## Abstract

**Background:** The quality of gene annotation determines the interpretation of results obtained in transcriptomic studies. The growing number of genome sequence information calls for experimental and computational pipelines for *de novo* transcriptome annotation. Ideally, gene and transcript models should be called from a limited set of key experimental data.

**Results:** We developed TranscriptomeReconstructoR, an R package which implements a pipeline for automated transcriptome annotation. It relies on integrating features from independent and complementary datasets: i) full-length RNA-seq for detection of splicing patterns and ii) high-throughput 5’ and 3’ tag sequencing data for accurate definition of gene borders. The pipeline can also take a nascent RNA-seq dataset to supplement the called gene model with transient transcripts.

We reconstructed *de novo* the transcriptional landscape of wild type *Arabidopsis thaliana* seedlings as a proof-of-principle. A comparison to the existing transcriptome annotations revealed that our gene model is more accurate and comprehensive than the two most commonly used community gene models, TAIR10 and Araport11. In particular, we identify thousands of transient transcripts missing from the existing annotations. Our new annotation promises to improve the quality of *A.thaliana* genome research.

**Conclusions:** Our proof-of-concept data suggest a cost-efficient strategy for rapid and accurate annotation of complex eukaryotic transcriptomes. We combine the choice of library preparation methods and sequencing platforms with the dedicated computational pipeline implemented in the TranscriptomeReconstructoR package. The pipeline only requires prior knowledge on the reference genomic DNA sequence, but not the transcriptome. The package seamlessly integrates with Bioconductor packages for downstream analysis.

## Background

Eukaryotic transcriptomes are large and complex: most genes can produce multiple isoforms, which may differ in their splicing pattern, localization of 5’ and 3’ ends and protein-coding potential [1]. RNA polymerase II continues well beyond the polyadenylation site as part of the mechanism of transcriptional termination, yet this read-through transcription is missing from gene models [2]. Moreover, transcripts generated from non-coding regions of the genome are often poorly annotated yet may exert regulatory functions even despite the absence of a stable RNA product [3–5]. Steady-state RNA sequencing methods offer little information on transient non-coding RNA species and read-through transcription. However, such transcription events result in overlapping transcription units which may contribute to gene expression regulation [6].

In the past, building a gene model for a species from EST libraries and cDNA clones sequenced by the Sanger method could require years of labor by an international consortium. Genome-wide detection of transcribed exons from RNA-seq data became feasible with short-read sequencing platforms such as Illumina. Indeed, transcript models for many less characterized genomes are primarily based on short-read RNA-seq data, e.g. *Pisum sativum* [7], *Oryza sativa* [8] and *Fragaria vesca* [9]. Even for the model plant species *Arabidopsis thaliana*, the commonly used transcriptome annotations TAIR10 and Araport11 are to a large degree based on Illumina RNA-seq datasets [10, 11].

However, the short read RNA-seq has some fundamental limitations. First, the RNA-seq coverage gradually decreases towards gene borders, which limits accurate definition of 5’ and 3’ ends of transcripts [12]. Second, although the positions of splice sites can be determined with high accuracy, the correct resolution of isoform splicing patterns is limited. Third, sequencing of steady-state RNA in wild type samples gives little information about transient non-coding transcripts. Taken together, these considerations offer a cautionary tale for gene annotations based on RNA-seq data.

Third generation sequencing techniques from Oxford Nanopore (ONT) and PacBio recently revolutionized the field of transcriptomics. Theoretically, a long RNA-seq read may cover the whole isoform from start to end, directly informing on the exon structure and splicing patterns. ONT also allows for direct sequencing of RNA molecules, thus eliminating biases associated with cDNA synthesis. This attractive feature makes ONT Direct RNA-seq a promising choice for *de novo* characterization of novel transcriptomes and validation of existing transcript models.

However, ONT Direct RNA-seq has four key limitations. First, up to 30-40% of bases can be called with errors [13, 14]. To tolerate the sequencing errors, the dedicated aligners allow for more mismatches and thus inevitably sacrifice the accuracy of alignments. As a result, the alignment software may fail to detect true genomic origin of the read or correctly define the exon-intron borders. Second, a fraction of long reads cover only 3’ portions of the original mRNA molecule, for example due to either RNA fragmentation, or premature termination of the sequencing reaction. Even when an intact mRNA molecule is fully sequenced from 3’ end to 5’ end, the last base of the read usually aligns a few tens of nucleotides downstream from the true transcription start site (TSS) [15]. Third, ONT sequencing is prone to homopolymeric tract skipping [16], which may appear as short exitrons (exonic introns) in Direct RNA-seq data. Fourth, the existing full-length RNA-seq protocols rely on RNA poly(A) tails for sequencing library construction. Therefore, non-polyadenylated RNA species which include many long non-coding RNAs (lncRNAs) will not be detected. Taken together, multiple factors preclude accurate transcript and gene model calling solely from the long reads.

Complementary genomics methods circumvent many of these limitations. For example, CAGE-seq [17] and PAT-seq [18] can detect RNA 5’ and 3’ ends, respectively, with high resolution, while nascent RNA methods such as GRO-seq [19] or NET-seq [20] can detect transient transcripts. Therefore, we posit that the highest quality transcript models can be constructed using integrative analysis levering the complementary strength of each of these methods.

We developed a *de novo* gene and transcript model construction pipeline TranscriptomeReconstructoR which takes three datasets as input: i) full-length RNA-seq (e.g. ONT Direct RNA-seq) to resolve splicing patterns; ii) 5’ tag sequencing (e.g. CAGE-seq) to detect TSS; iii) 3’ tag sequencing (e.g. PAT-seq) to detect polyadenylation sites (PAS). Optionally, it can also take a nascent RNA-seq dataset (e.g. NET-seq) to find transient RNAs. The pipeline returns the discovered genes and transcripts. We also included the option to refine the *de novo* gene and transcript models by the existing transcriptome annotation. TransctiptomeReconstructoR thus can be used for validation or improvement of existing gene models, as well as for data-driven annotation of non-model species with no prior gene model available.

## Implementation

We implemented the pipeline as an R package, available from the dedicated repository on GitHub (https://github.com/Maxim-Ivanov/TranscriptomeReconstructoR). The package is based on Bioconductor packages (GenomicRanges, GenomicAlignments, rtracklayer), as well as on tidyverse and collections packages from CRAN [21, 22]. It takes aligned BAM files as input and returns a set of GRanges and GRangesList objects which represent the gene and transcript models. These GenomicRanges objects can be either exported as BED files for visualization in genomic browsers, or directly used as input for downstream analysis by various packages available from the Bioconductor. The TranscriptomeReconstructoR workflow is streamlined and includes 6 consecutive function calls (optionally 8, if the nascent RNA-seq dataset is used). The accompanying vignette contains the user manual, as well as in-depth description of the algorithm (Additional File 1).

The underlying basis for TranscriptomeReconstructoR is the correction and validation of long RNA-seq reads by the positions of TSS and PAS that are called from the independent short read 5’ and 3’ tag datasets. Long reads from ONT Direct RNA-seq or PacBio Iso-Seq are extended towards nearby TSS and/or PAS, given that the extension distance does not exceed the reasonable limit (100 bp by default). After the extension, the long reads are classified as “complete” or “truncated”, depending on the overlap with the independently called TSS and PAS (Fig. 1A). The extension procedure allows to rescue a substantial fraction of long reads and simultaneously decrease the number of artifact transcript isoforms with alternative 5’- or 3’ terminal exons.

**Figure 1.**
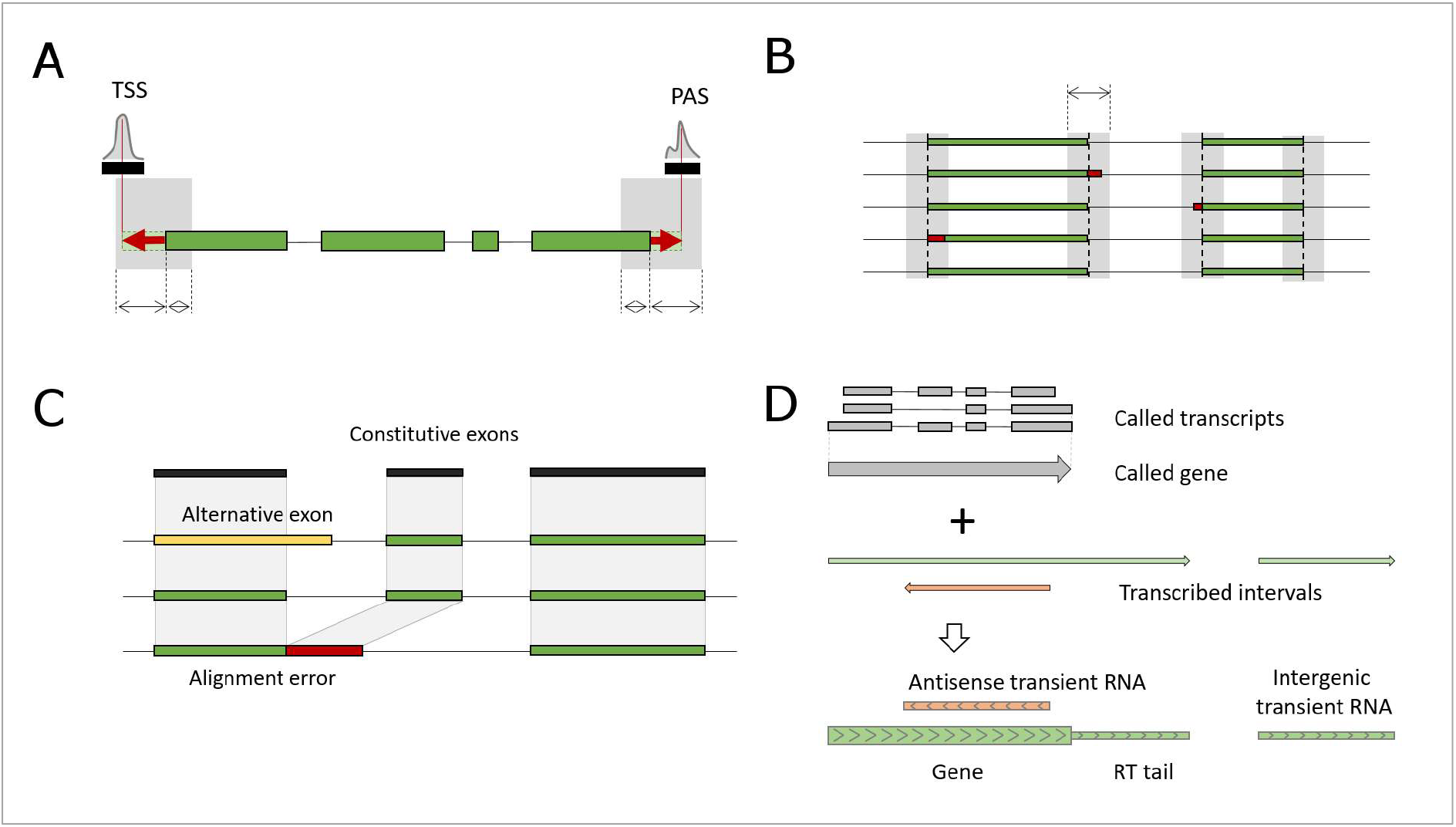
Outline of the TranscriptomeReconstructoR concept. **(A)** Genomic coordinates of TSS and PAS are called from 5’ and 3’ tag sequencing data, respectively. Terminal subalignments of long reads are extended towards the summits of the nearby TSS and PAS (within 100 bp distance on the same strand). An extended read is considered complete, if its 5’ and 3’ ends overlap with TSS and PAS, respectively. **(B)** Overlapping subalignments of complete reads are clustered together, if the pairwise distances between their borders do not exceed 10 bp. Within each cluster, the coordinates of subalignments are unified to the most frequently observed values. **(C)** Long reads sharing the same TSS and PAS are grouped together. Subalignments present in more than 50% of reads within the group are considered constitutive exons. Alignment errors are detected by comparing subalignments of each long read to the set of constitutive exons. A subalignment with alternative 3’ or 5’ border can be considered a novel alternative exons, only if the next subalignment in the same read precisely matches the next constitutive exon. Otherwise, if the next constitutive exon is absent from the read, the tested subalignment is marked as potential alignment error. **(D)** Continuous intervals of nascent transcription (green) cover a larger fraction of genome than the transcripts and genes called from steady-state long read RNA-seq data (grey). Nascent transcription intervals which do not overlap with regions of mature RNA production on the same strand, are classified into either read-through (RT) tails, or transient RNAs.

The method also suppresses the alignment noise of long reads by the adjustment of 5’- and 3’ splice sites. The true alternative splice sites are assumed to be separated by a certain minimal distance (10 bp by default). Exonic subalignments of long reads with 5’ or 3’ borders differing by less than this value are grouped together, and their coordinates are unified by the majority vote (Fig. 1B). Otherwise the “fuzzy” borders of subalignments might inflate the number of alternative 5’ and 3’ splicing events.

A common problem with full-length RNA sequencing reads aligned by dedicated aligners such as Minimap2 [23] is the under-splitting, i.e. erroneous extension of an exon over the adjacent intronic region [24]. As a result, the next exon appears as missing from such read, although it is present in other reads aligned within the same locus. The most probable explanation for this phenomenon is the inherent low sensitivity of long read aligners [24]. Assuming that the majority of reads still align correctly, we detect such alignment errors by comparing each subalignment in a long reads to the linear sequence of constitutive exons, that we extract from the whole set of long reads aligned to given locus. Subalignment with an alternative 5’- and/or 3-border relative to a constitutive exon are considered valid alternative exons, only if the next constitutive exon is also present in the read (Fig. 1C). Otherwise, the alignment is marked as a possible alignment error. Reads containing at least one alignment error are skipped from the transcript calling procedure, thus further decreasing the number of artifact alternative 5’ and 3’ splice sites.

Finally, identical long reads without alignment errors are collapsed into transcripts. Such transcripts are divided into High Confidence (HC), Medium Confidence (MC) and Low Confidence (LC) groups, depending on the support from TSS and PAS datasets. HC transcripts are constructed from long reads which start in a TSS and end in a PAS (i.e. are supported by all three datasets). MC transcripts have either TSS or PAS, whereas LC transcripts are supported by long reads only. The MC and LC transcripts may originate from partially fragmented RNA molecules and thus may have unreliable outer borders. They are called to rescue the loci where no complete reads were discovered, perhaps due to low expression level. Furthermore, the called transcripts are clustered into HC, MC and LC genes.

If a nascent RNA-seq dataset is available, the gene model can be further improved by intervals of nascent transcription which can be found outside of the called gene boundaries (Fig. 1D). These often represent transient and/or non-polyadenylated RNAs which escape detection by the poly(A)-dependent steady-state RNA sequencing methods. Another phenomenon which can be observed only in the nascent RNA-seq track is read-through transcription. The read-through (RT) “tails” immediately downstream from the protein-coding genes are explained by the “torpedo” termination model where RNAPII elongation may continue for up to a few Kb beyond the cleavage and polyadenylation site [2]. If such “tails” were identified, we appended them to the respective called gene.

TranscriptomeReconstructoR returns a set of GRanges and GRangesList objects which exhaustively annotate the transcriptome. They contain the following information: i) The coordinates of HC, MC and LC genes; ii) The length of RT tails; iii) The exon-intron structure of HC, MC and LC transcripts; iv) The coordinates of intergenic and antisense transient RNA; v) The exon-intron structure of fusion transcripts (i.e. transcripts which cover two or more adjacent genes due to inefficient termination). These GenomicRanges objects can be easily exported as BED files for visualization in genomic browsers.

## Results

### *De novo* annotation of the Arabidopsis transcriptome

We tested and validated TranscriptomeReconstructoR on 2 weeks old wild type *Arabidopsis thaliana* seedlings (Col-0 ecotype). The following published datasets were used as input: ONT Direct RNA-seq [25], CAGE-seq [26], PAT-seq [27] and plaNET-seq [28]. The CAGE-seq and PAT-seq data produced 30973 TSS and 36729 PAS, respectively. Using these data, out of 3.7M raw ONT reads, 43.8% started in a TSS, 78.6% ended in a PAS, and 35.2% satisfied both conditions. After extension of 5’ and 3’ ends of the long reads towards nearby TSS and PAS within 100 bp (see Fig. 1A), the fraction of “complete” reads was more than doubled, raising from 35.2% to 71.6%. The extended long reads contained 14.5M exonic subalignments. Among them, 531,690 internal subalignments were adjusted at either 5’- or 3’ splice site by the majority vote within 10 bp offset (see Fig. 1B). As a result, the number of unique exons has decreased by 9.6% (from 965,012 to 872,565). Thus, small alignment errors at exon borders may inflate the diversity of called exons. In addition, 3.5% of the long reads likely contained at least one alignment error (see Fig. 1C). These results demonstrate that the complexity of alternative isoforms could be substantially overestimated, if long reads were neither validated by the orthogonal 5’ and 3’ tag sequencing data nor corrected by comparison to the common alignment pattern of the locus.

This final set of corrected long reads was used to call HC, MC or LC transcripts which were further clustered into genes. Minor isoforms (supported by less than 1% of long reads in given gene) were skipped. The final annotation consists of 65864 HC transcripts in 15884 HC genes, 4914 MC transcripts in 3478 MC genes and 2092 LC transcripts in 1799 LC genes. In addition, we detected 438 fusion transcripts that span two or more adjacent genes.

As final step, we augmented the gene model with nascent transcription data from the plaNET-seq dataset (see Fig. 1D). We found that 86.5% of all called genes have a read-through (RT) tail longer than 50 bp, and the median length of such RT tails was 351 nt. In addition, we found 5064 transient transcripts that are supported only by plaNET-seq reads and likely correspond to unstable and/or non-polyadenylated lncRNAs. The *de novo* annotation for *A. thaliana* is available from author’s GitHub (see Materials and methods).

### Validation of the *de novo* annotation

The gene and transcript models generated by TranscriptomeReconstructoR were compared to the two existing annotation sets for Arabidopsis - TAIR10 and Araport11 [10, 11]. It is important to note that these two gene annotation builds are markedly different. The TAIR10 is conservative, focused on protein-coding genes and contains very few non-coding transcripts. Conversely, Araport11 includes thousands of non-coding RNA genes not present in TAIR10. In addition, gene borders are substantially different between these two annotations. In particular, Araport11 genes tend to be longer, i.e. they have more upstream 5’ ends and more downstream 3’ ends compared to their mates in TAIR10 (Fig. S1 in Additional File 2). Since our pipeline is agnostic to current annotations and used independent datasets, we reasoned that the resulting output could help to assess the accuracy of existing annotations. To this end, we compared our *de novo* annotation to TAIR10 and Araport11 at the gene and exon levels.

In general, our HC and MC genes agreed well with known genes from both annotations. In particular, more than 95% of HC genes have a unique mate (i.e. overlap with a single annotated gene on the same strand) in both TAIR10 and Araport11 (Fig. 2A). The majority of called genes have strong base-pair overlap (defined as intersection to union ratio of distances between 5’ and 3’ gene borders) with their mates in TAIR10 and Araport11 (Fig. 2B). For example, 85% and 61% of HC genes had at least 90% overlap with TAIR10 and Araport11 genes, respectively. Moreover, 5’ and 3’ borders of the called genes often coincided with respective borders of TAIR10 genes (Fig. 2C, upper panel).

**Figure 2.**
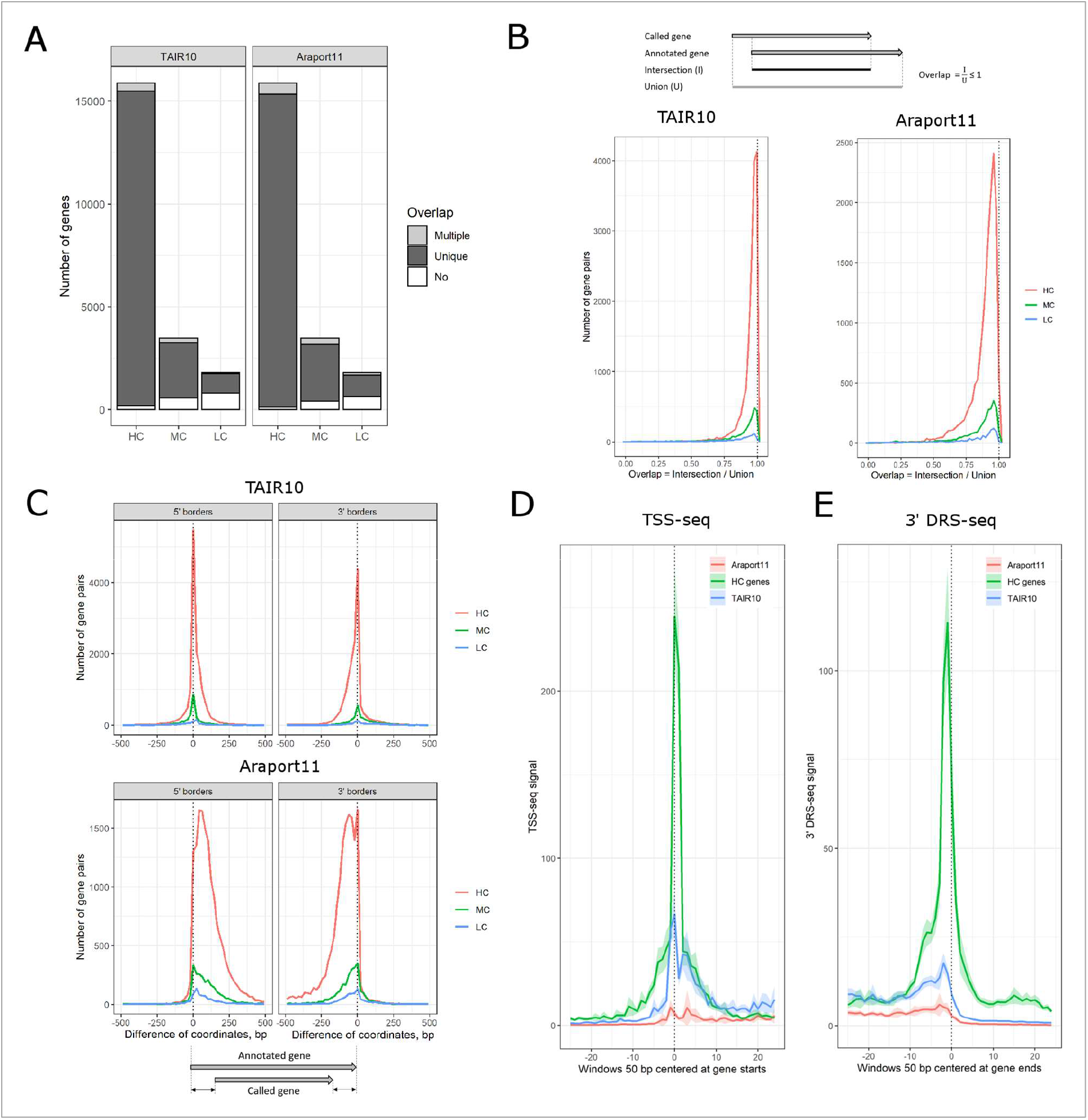
Comparison of called gene borders to TAIR10 and Araport11. **(A)** Stacked barplot shows the counts of High Confidence (HC), Medium Confidence (MC) and Low Confidence (LC) genes which either: a) have no overlap with any annotated gene (white); b) have a unique mate in the annotation (i.e. overlap a single annotated gene which in turn does not have any other overlaps among the called genes; dark grey); c) have multiple matches in the annotation (light grey). Only overlaps on the same strand are considered valid. Only the uniquely matched pairs of genes were used on the next subfigures. **(B)** Distribution of pairwise overlap values between called HC (red), MC (green) or LC genes (blue) and their unique mates in TAIR10 or Araport11. Y axis shows the number of gene pairs, X axis shows the overlap calculated as the ratio of intersection (common length) to union (total length) of the overlapping genomic intervals. Area under the curve is proportional to the number of gene pairs in the group. **(C)** Distribution of differences between 5’ or 3’ borders (left and right panels, respectively) of the matched gene pairs. Y axis shows the number of gene pairs, X axis shows the difference of genomic coordinates (in bp). A negative (positive) difference value means that the border of the called gene is located upstream (downstream) from the respective border of its mate in TAIR10 (upper panel) or Araport11 (lower panel). A narrow peak with summit at zero position on the X axis means that the gene pairs most often have identical positions of the borders. A smooth peak with multiple summits indicates high incidence of mismatched gene borders. Area under the curve is proportional to the number of gene pairs in the group. **(D)** Metagene profile of TSS-seq signal over 5’ gene borders. The HC/TAIR10 and HC/Araport11 matched gene pairs were joined into HC/TAIR10/Araport11 triads. Thus, each gene has three alternative 5’ borders predicted by TranscriptomeReconstructoR, TAIR10 and Araport11. Fixed length genomic intervals (50 bp) were centered on the 5’ gene borders in each of the three groups. TSS-seq signal (which is proportional to TSS usage) was averaged among the genomic windows. Y axis shows the average sequencing coverage, X axis shows the genomic coordinates relative to the 5’ gene border (zero corresponds to the predicted gene start). The color of the wiggle line indicates the origin of the genomic windows: blue for TAIR10, red for Araport11 and green for the called HC genes. **(E)** Metagene profile of Helicos 3’DRS-seq signal over 3’ gene borders. Both TSS-seq (in panel D) and 3’ DRS-seq (in panel E) demonstrate a sharp peak at the respective gene borders derived from TAIR10 and HC genes, but not from Araport11 genes. This indicates that gene borders predicted by Araport11 often disagree with the experimental evidence. Moreover, the HC peak (green) is substantially higher than the TAIR10 peak (blue) in both TSS-seq and 3’ DRS-seq. This means that TSS and PAS positions predicted from the HC gene set are in a better agreement with the experimental data than the TAIR10.

On the other hand, the gene borders may differ between called genes and their mates in the existing annotations. Notably, the called 5’ and 3’ gene borders were systematically shifted downstream and upstream, respectively, from the genomic positions predicted by Araport11 (Fig. 2C, lower panel). This is consistent with the observation that Araport11 genes are in general wider than in TAIR10 (see Fig. S1 in Additional File 2). Moreover, we found that 5’ borders of HC genes in our gene model are in a better agreement with independent experimental mapping of 5’ boundaries by TSS-seq [29] than positions predicted from either TAIR10 or Araport11 (Fig. 2D). Similarly, 3’ borders of HC genes coincided with Helicos 3’ DRS-seq [30], an independent method to detect PAS, substantially better than 3’ ends of the corresponding annotated genes (Fig. 2E). Transcription of *Arabidopsis* genes frequently produces short promoter-proximal RNAs (sppRNA) which are sensitive to the *HEN2* exonuclease and terminate about 100 nt downstream from the TSS [31]. We found that 5’ borders of genes called by TranscriptomeReconstructoR are in a better agreement with the sppRNA termination sites than 5’ borders of their annotated mates in both TAIR10 and Araport11 (Fig. S2 in the Additional File 2). This finding supports the idea that the gene models resulting from TrancriptomeReconstructoR are in better agreement with independent transcriptomic datasets than both commonly used annotations.

We also analyzed the base-pair overlaps of internal exons between the called genes and their mates in each of the existing annotations. For more than 78% of called internal exons there was a unique known matching exon (where both 5’ and 3’ borders differed by no more than 5 bp) when compared to either TAIR10 or Araport11 (Fig. 3A). Within those, the vast majority of splicing acceptor and donor sites were exact matches (Fig. 3B). The remaining 22% called exons could represent novel alternative exons. The most prominent class among them are novel intron retention (IR) events, whereas alternative 5’ or 3’ exon borders are less common (see Fig. 3A).

**Figure 3.**
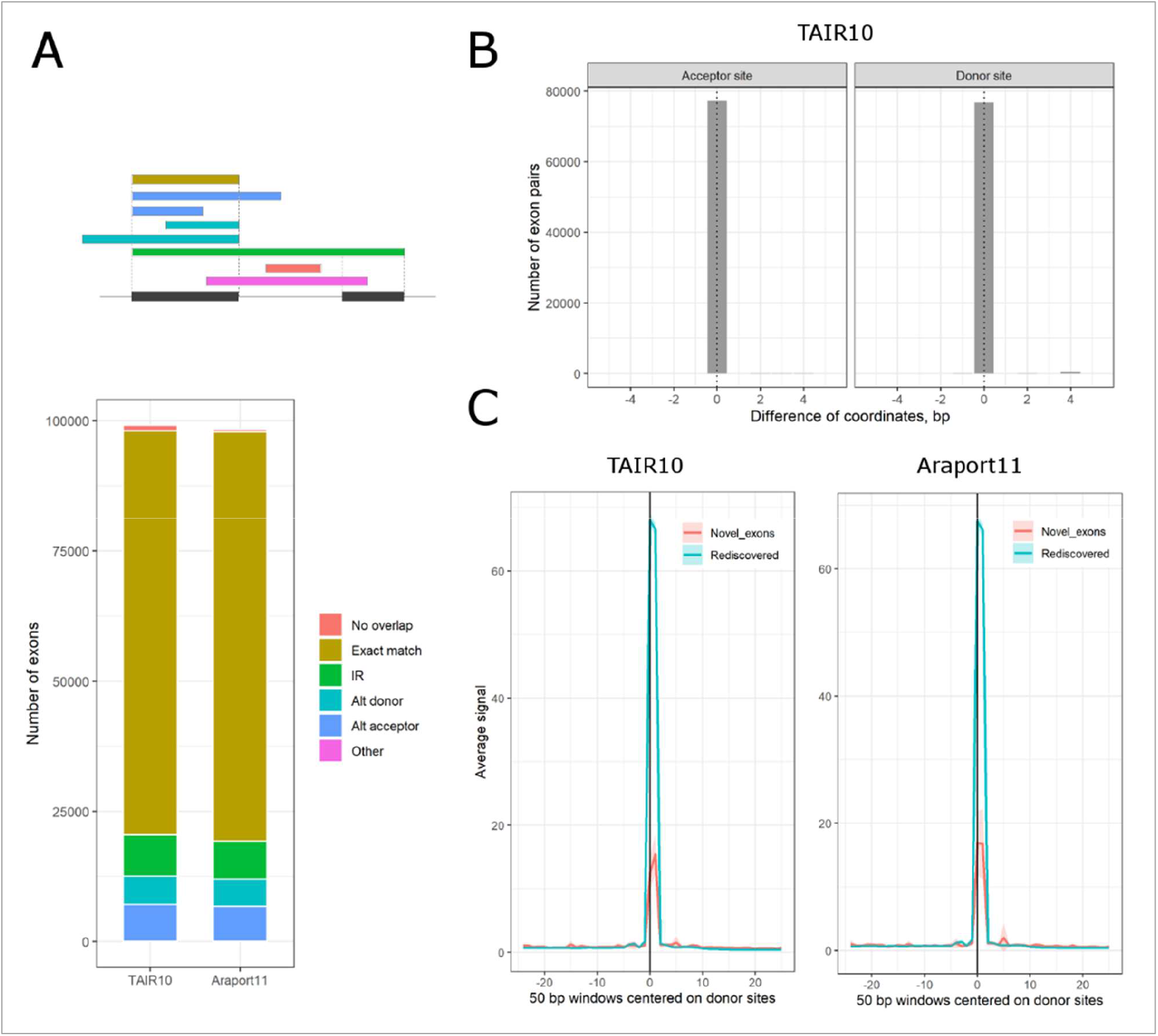
Comparison of called exon borders to TAIR10 and Araport11. **(A)** Overlaps of called internal exons vs TAIR10 and Araport11. On top, the schematic outline shows the possible overlap types for internal exons: i) exact match (within 5 bp offset) at both borders of the same annotated exon; ii) exact match at borders of two adjacent exons (intron retention); iii) exact match only at 5’ exon border (alternative acceptor site); iv) exact match only at 3’ exon border (alternative donor site). Notably, it is also possible that a called exon has no overlap with any annotated exon, or that the overlapping pattern is more complicated than those shown on the schematic. The bottom stacked barplot shows the counts of called internal exons with different type of pairwise overlaps, colored as in the schematic above, with their best mate exons in either TAIR10 or Araport11. **(B)** Histogram shows the distribution of called 5’ or 3’ exon borders (left and right panel, respectively) relative to the acceptor and donor sites from their best mates in TAIR10. The central (zero) position on X-axis corresponds to exact match between called and annotated exon borders. **(C)** Metagene plot shows the average density of spliceosome intermediate reads in the plaNET-seq dataset over 50 bp windows centered at 3’ exon borders (donor splice sites) of either re-discovered (blue) or newly discovered (red) called exons. The shaded areas show the normal-based confidence intervals for the mean. The presence of sharp peaks exactly at the called 3’ exon borders (at zero position on the X axis) in the newly discovered exons indicate that they represent functional donor sites. The absolute peak height is proportional to the average expression levels of genes in each group.

We also found 740 and 337 exons which do not overlap with any known exon in TAIR10 and Araport11, respectively. These could represent newly discovered exons. To test this hypothesis, we used the plaNET-seq dataset. We demonstrated that the raw plaNET-seq data are enriched for reads with the first base aligned exactly to the donor splice sites [28]. These reads correspond to splicing intermediates, and they appear in the plaNET-seq data due to co-purification of the spliceosome together with the transcriptionally engaged RNAPII complex. Interestingly, we found that such splicing intermediate reads are also enriched at the newly discovered donor sites (Fig. 3C). Thus, the plaNET-seq data support the conclusion that TranscriptomeReconstructoR discovered *bona fide* new exons.

Our annotation also identified novel genes. We found 63 HC genes, 234 MC genes, 428 LC genes and 2094 transient transcripts that did not overlap on the same strand with any known gene from either TAIR10, Araport11 or the custom lncRNA catalogue defined by Zhao and co-authors [32]. However, the vast majority of these novel transcription units (TUs) were found antisense to known genes in TAIR10 and/or Araport11 (Fig. 4A). The novel TUs were validated by independent pNET-seq dataset which shows the intensity of nascent RNAPII transcription [33]. The averaged pNET-seq signal is markedly increased over the bodies of the novel TUs and decreased at their borders, thus confirming their transcriptional activity (Fig. 4B). Importantly, transposons offer no explanation for these novel TUs. Only 1 MC gene, 2 LC genes and 14 transient RNA have at least 50% overlap with any transposon discovered in the comprehensive study of Panda and co-authors [34].

**Figure 4.**
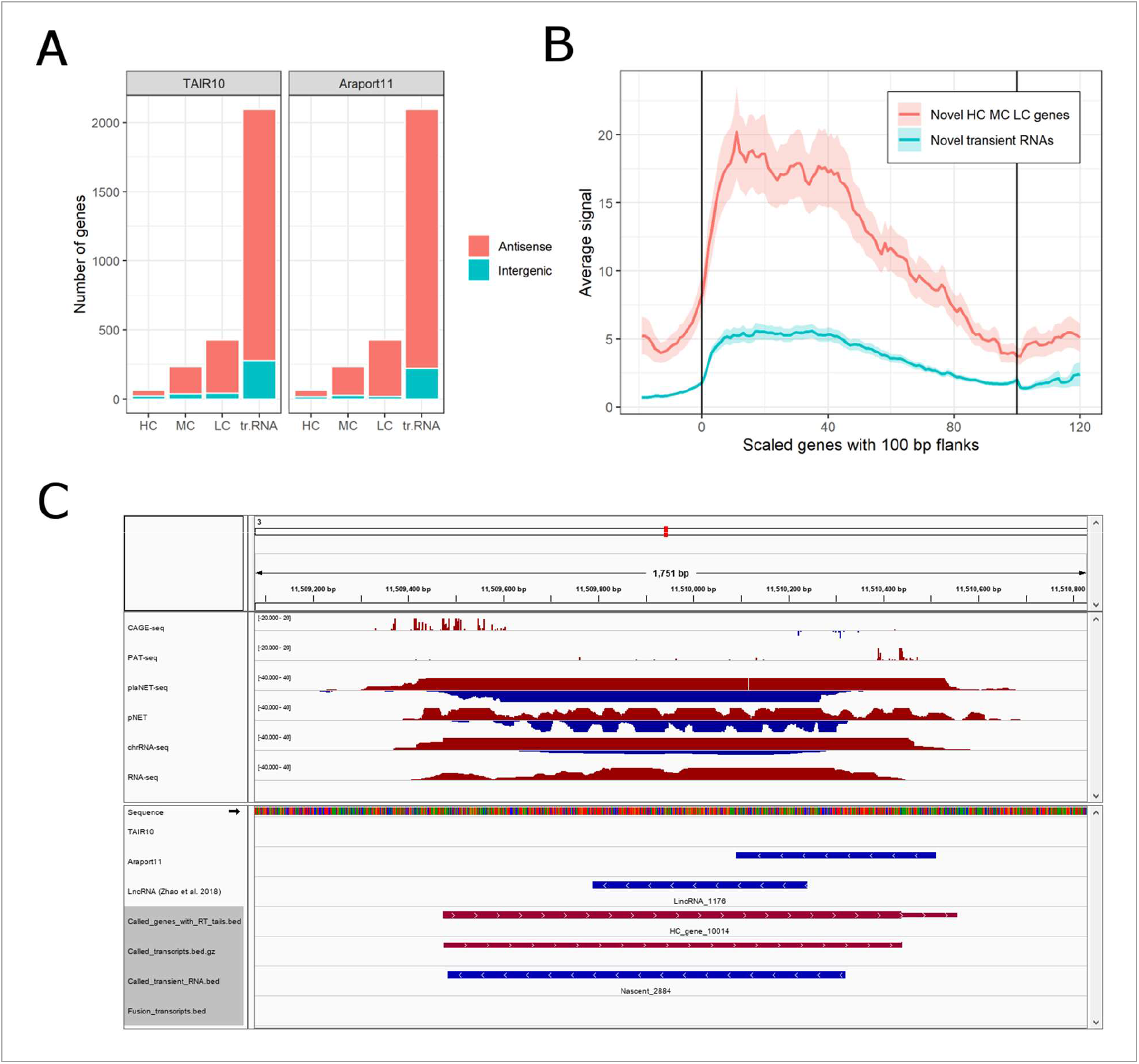
Novel genes and transient RNAs. **(A)** Stacked barplot shows the counts of novel HC, MC and LC genes and transient RNAs which have no overlap with any known gene or lncRNA on the same strand. Novel transcription units which overlap any known gene on the opposite strand are considered antisense (red), otherwise intergenic (blue). **(B)** Metagene plot of pNET-seq signal over the whole bodies of novel HC, MC and LC genes and novel transient RNAs. The called genes have variable length, therefore they were scaled to 100 bins. The 100 bp upstream and downstream flanking regions were scaled to 20 bins each. The vertical lines in the plotting area denote the starts and ends of novel genes. Red and blue wiggle lines show the average RNAPII elongation activity in novel genes and transient RNAs, respectively. Red and blue shaded areas show normal-bases 95% confidence interval for the respective means. **(C)** Example of a novel gene encoding a stable transcript. Features on forward and reverse strands are shown in red and blue, respectively. HC_gene_10019 is a High Confidence gene which was called on forward strand, i.e. in antisense orientation to lncRNA-encoding locus (denoted as AT3G05945 in Araport11, and as LincRNA_1146 in the annotation of Zhao *et al*., 2018). This novel gene has support from both Nanopore Direct RNA-seq (not shown), PAT-seq, CAGE-seq and plaNET-seq. Since the first two methods depend on the presence of poly(A) tail, the transcript is most probably polyadenylated. Moreover, the gene was validated by three independent datasets (pNET-seq, chrRNA-seq and RNA-seq). Given that the gene is clearly visible even in RNA-seq data, it remains unclear why it is absent from both TAIR10 and Araport11 annotations.

Representative screenshot of a novel HC gene (Fig. 4C) shows that it is further supported by two other independent datasets - chrRNA-seq and stranded RNA-seq [35, 36]. These two complementary methods show nascent RNA or mature RNA, respectively. In contrast, a representative novel transient RNA (Fig. S3 in Additional File 2) is supported by chrRNA-seq signal but not by RNA-seq, consistent with a transcript subject to nuclear RNA degradation.

Our gene model also resolves potential mis-annotations in TAIR10 and Araport11. For example, a gene may appear as a continuous transcription unit in the existing annotations, whereas our results suggest that it consists of two or more independent non-overlapping transcripts. We detected 82 TAIR10 genes which overlap two or more called genes by more than 75% of their lengths on the same strand. A representative example of such gene on Fig. S4 (see Additional File 2) demonstrates that its split nature can be further supported by independent TIF-seq dataset which detects both ends of mature RNA transcripts [31].

## Discussion

We developed TranscriptomeReconstructoR motivated by the observation that Araport11, the *A.thaliana* annotation considered to be the most comprehensive in terms of non-coding transcripts, has highly inaccurate TSS positions [26]. This inaccuracy limits genome-wide analyses that are dependent on accurate positions of gene promoters. For example, analyses for enriched sequence motifs at a fixed distance from mis-annotated TSS may lead to incorrect results. Previously, we demonstrated low sensitivity of metagene profile analysis at 5’ and 3’ gene borders predicted from Araport11, compared to TAIR10 [28]. Conversely, the “classical” TAIR10 annotation is too conservative, as it is focused on protein-coding genes and largely ignores non-coding transcription. However, the *Arabidopsis* genome is rich in non-coding transcripts [37]. Thus, there is no “gold standard” gene annotation, even for a well-studied model species as *A.thaliana*. We built a new annotation to test if a data-driven gene model could reflect the actual transcriptional landscape of wild type *A.thaliana* seedlings more accurately than both TAIR10 and Araport11. Refining an existing annotation with experimental data offers one possible solution. While this approach can be useful [28], errors and limitations of the input gene model would be inherited by the output annotation. To avoid this problem, we focused on a *de novo* transcriptome reconstruction approach.

We tested the performance of TranscriptomeReconstructoR for fully automated calling of gene and transcript models on four previously published datasets corresponding to 2 weeks old *A.thaliana* seedlings. Impressively, our *de novo* gene model could outperform both TAIR10 and Araport11 in certain aspects, for example in determining the 5’ and 3’ gene borders (see Fig. 2D-E and Fig. S2). Thus, an important advantage of our annotation is the improved accuracy and sensitivity of metagene profile analysis around TSS and PAS. Moreover, we resolved some historical errors of TAIR10 and Araport11, as exemplified by two closely spaced genes which were merged into a single gene in both annotations (see Fig. S4 in Additional File 2). In addition, TranscriptomeReconstructoR detected more than two thousand novel transcripts, most often in antisense orientation to the known genes, that were completely absent from the existing annotations (see Fig. 4A). Taken together, these observations validate the approach implemented in TranscriptomeReconstructoR and suggest that the new annotation can complement and enhance the TAIR10 and Araport11 gene models in *A. thaliana* transcriptomic studies.

The list of newly sequenced species from all kingdoms of life is rapidly growing. At present, a popular way of constructing transcriptomes of the emerging species is the transcript assembly from the short read RNA-seq data. TranscriptomeReconstructoR offers an attractive alternative with improved accuracy. We anticipate that research groups, rather than consortia, will employ this strategy to characterize draft transcriptomes in non-model species with recently assembled genomes.

## Conclusions

TranscriptomeReconstructoR is a user-friendly tool that combines Next generation and Third generation sequencing datasets for *de novo* calling of gene and transcript models. It offers assembly of draft transcriptome annotations in recently sequenced non-model species.

## Materials and methods

Only publicly available datasets were used in this study. Direct RNA-seq and PAT-seq data were downloaded from the European Nucleotide Archive (accession numbers PRJEB32782 and SRP145554, respectively), Helicos 3’ DRS-seq data from the DNA Data Bank of Japan (accession ERP003245), all other datasets from the NCBI Sequence Read Archive: CAGE-seq (GSE136356), plaNET-seq (GSE131733), pNET-seq (GSE109974), TSS-seq (GSE113677), chrRNA-seq (PRJNA591665), RNA-seq (GSE81202) and TIF-seq (GSE129523). Supplementary code to process the raw data and completely reproduce the results obtained in this study is available from https://github.com/Maxim-Ivanov/Ivanov_et_al_2021. The data processing pipeline requires the TranscriptomeReconstructoR package which can be installed from https://github.com/Maxim-Ivanov/TranscriptomeReconstructoR. The new annotation for *A.thaliana* was deposited on https://github.com/Maxim-Ivanov/Ivanov_et_al_2021/tree/main/Annotation.

## Supporting information

Vignette to TranscriptomeReconstructoR

## Additional files

**File name:** Additional File 1

**File format:** PDF

**Title of data:** Vignette to TranscriptomeReconstructoR.

**Description of data:** The file describes an example pipeline for *de novo* calling of gene and transcript models by TranscriptomeReconstructoR, as well as detailed description of the algorithm.

**File name:** Additional File 2

**File format:** PDF

**Title of data:** Supplementary Figures S1-S4.

**Description of data:**

**Figure S1** (Comparison of gene borders between TAIR10 and Araport11). Pairs of TAIR10 and Araport11 genes were matched by gene identifiers. The frequency polygon plot shows the distribution of differences (in bp) between 5’ or 3’ borders (left and right panels, respectively) of the matched gene pairs. A negative difference value means that the gene has a more upstream coordinate in Araport11 than in TAIR10. Similarly, a positive value means that the coordinate is downstream in Araport11 relative to TAIR10. The plot shows that Araport11 genes often start upstream and end downstream from their mates in TAIR10, i.e. are in general wider.

**Figure S2** (Metagene plot of PAS signal around TSS of sppRNA-containing genes). Triads of called genes, TAIR10 and Araport11 genes were matched by overlaps of genomic coordinates. The triads were subsetted to sppRNA-containing genes (n = 1326). Genomic windows (600 bp) were produced around the TSS positions. TIF-seq reads from the exosome mutant *hen2-2* sample were resized to their last bases (which correspond to the empirical PAS sites). The PAS signal was plotted along each of the 3 sets of genomic windows, and the metagene plots were overlaid. The vertical line corresponds to the TSS position. The average PAS signal plotted in TSS-centered windows predicted by the called genes (green) forms the most sharp peak at 100 bp downstream from the TSS. This means that 5’ borders of the called genes are in a better agreement with the PAS sites of sppRNAs than 5’ borders of the same genes in the existing annotations.

**Figure S3** (Example of a novel gene encoding transient RNA). Features on forward and reverse strands are shown in red and blue, respectively. The called transient RNA dubbed “Nascent_0037” is encoded on the forward strand. i.e. in antisense orientation to the AT1G05010 gene (ethylene-forming enzyme). This novel RNA transcript has no support from Nanopore Direct RNA-seq data (not shown). It was called from plaNET-seq data only, thus suggesting its transient and/or non-polyadenylated nature. The existence of this transcript was validated by two independent datasets (pNET-seq and chrRNA-seq) which are enriched for nascent RNA species. As expected, it is not visible in RNA-seq dataset, which explains why it is lacking from both TAIR10 and Araport11.

**Figure S4** (Example of a gene misannotated in TAIR10 and Araport11). Features on forward and reverse strands are shown in red and blue, respectively. The AT3G20270 (AtLBR-2) gene involved in pathogene response is annotated as a single gene in both TAIR10 and Araport11. However, TranscriptomeReconstructoR shows that it consists of two non-overlapping genes in tandem orientation. Both of these genes were classified as High Confidence, because they were supported by Nanopore Direct RNA-seq (not shown), CAGE-seq and PAT-seq data. The presence of two individual transcripts was further validated by independent TIF-seq data (the lowest track).

## Abbreviations

RNA-seq: RNA sequencing;
ONT: Oxford Nanopore;
PacBio: Pacific Biosciences;
NET-seq: Native Elongation Transcript sequencing;
GRO-seq: Global Run-On sequencing;
A. thaliana: Arabidopsis thaliana;
cDNA: complementary DNA;
TSS: transcription start site;
PAS: polyadenylation site;
lncRNA: long non-coding RNA;
CAGE-seq: Cap Analysis of Gene Expression sequencing;
PAT-seq: Poly(A) tag sequencing;
CRAN: Comprehensive R Archive;
BAM: Binary Alignment Map;
BED: Browser Extensible Data;
Iso-seq: isoform seqiencing;
HC: high confidence;
MC: medium confidence;
LC: low confidence;
RT: read-through;
plaNET-seq: plant Native Elongation Transcript sequencing;
M: million;
bp: base pair;
ncRNA: non-coding RNA;
TSS-seq: Transcription Start Site sequencing;
3’ DRS-seq: 3’ Direct RNA sequencing;
TU: transcription unit;
chrRNA-seq: chromatin-associated RNA sequencing;
TIF-seq: Transcript Isoform sequencing;
EST: expressed sequence tag;

## Acknowledgements

We would like to acknowledge all members of the Marquardt’s and Sandelin’s labs for discussion on the algorithm and useful feedback on the new annotation.

## Authors’ contribution

MI conceived the project and implemented the code. MI, AS and SM wrote the manuscript. All authors read and approved the final manuscript.

## Availability of data and materials

The TranscriptomeReconstructoR package if freely available from GitHub:

**Project name:** TranscriptomeReconstructoR

**Project homepage:** https://github.com/Maxim-Ivanov/TranscriptomeReconstructoR

**Operating system(s):** Platform independent

**Programming language:** R

**Other requirements:** CRAN, Bioconductor

**Licence:** GPL-3

**Any restrictions to use by non-academics:** GPL-3

All analyses are based on publicly available datasets obtained from GEO (for accession numbers, see Materials and methods). The *de novo* annotation for *A.thaliana* is available from https://github.com/Maxim-Ivanov/Ivanov_et_al_2021/tree/main/Annotation.

## Ethics approval and consent to participate

Not applicable.

## Consent for publication

Not applicable.

