## Supplementary material for "TrancriptomeReconstructoR: data-driven annotation of complex transcriptomes": Vignette to TranscriptomeReconstructoR

**Maxim Ivanov**

**December 04, 2020**

### Introduction

The *TranscriptomeReconstructoR* package implements a pipeline which allows for *de novo* assembly of transcriptomes by combining data from a few mutually orthogonal datasets:

1. Stranded full-length RNA sequencing (e.g. ONT Direct RNA-seq or PacBio Iso-Seq);
2. 5’-tag RNA sequencing (e.g. CAGE-seq or TSS-seq);
3. 3’-tag sequencing (e.g. PAT-seq or 3’READS);
4. (optionally) Nascent RNA-seq (e.g. NET-seq, GRO-seq or chrRNA-seq).

Observe that all input datasets are expected to be stranded, i.e. strand orientation of a read must correspond to strand orientation of the original RNA molecule. This means that certain library preparation protocols for Third generation RNA sequencing may be not compatible with the pipeline (or may require an additional pre-processing step to recover the strand information). For more details on the various full-length RNA-seq methods, see **Appendix 1**.

The full-length RNA-seq provides information on exon-intron structure of the original RNA transcripts. However, the long reads from Third generation sequencing platforms (both ONT and PacBio) suffer from certain biases, in particular high error rate and truncated 5’-ends. The high error rate inevitably results in lower accuracy of alignments. Therefore, a long read cannot be interpreted as a *bona fide* transcript isoform.

*TranscriptomeReconstructoR* aims to compensate for sequencing and alignment errors of long reads by:

- Extracting common features from multiple long reads originating from the same isoform;
- Correcting truncated 5’- and 3’-end of long reads by TSS and PAS coordinates which are derived from independent 5’- and 3’-tag sequencing datasets, respectively;

The pipeline returns gene and transcript models which are called from experimental data and thus are agnostic to any existing annotation. However, the resultant *de novo* transcriptome assembly may be refined by an existing annotation, if desired (see **Appendix 2**). In addition, *TranscriptomeReconstructoR* can supplement the transcriptome assembly by transient and/or non-polyadenylated transcripts (given that a nascent RNA-seq dataset was included).

*TranscriptomeReconstructoR* can be used to assemble draft transcriptomes of newly sequenced organisms, as well as to validate the existing gene annotations.

### Installation

The easiest way to install *TranscriptomeReconstructoR* is to use the `devtools::install_github()` function:

```
if (!"devtools" %in% rownames(installed.packages())) {
  install.packages("devtools")
}
devtools::install_github("Maxim-Ivanov/TranscriptomeReconstructoR",
                         build_vignettes = TRUE)
```

Then attach the installed package:

```
library(TranscriptomeReconstructoR)
```

### Input files

*TranscriptomeReconstructoR* uses BAM files as input. In this tutorial, we will use example BAM files obtained from the following published datasets of 2 weeks old *Arabidopsis thaliana* seedlings:

- Direct RNA-seq by ONT from Parker et al. (2020);
- CAGE-seq from Thieffry et al. (2020);
- PAT-seq from Yu, Lin, and Li (2019);
- plaNET-seq from Kindgren, Ivanov, and Marquardt (2020).

Details on downloading, pre-processing and alignment of the raw sequencing data can be found here. The BAM files were subsetted to a 300Kb interval on chromosome 1 (1:10,000,000-10,300,000). To retrieve the filenames, run the following code:

```
pkg <- "TranscriptomeReconstructoR"
drs_bamfiles <- system.file("extdata",
                            paste0("ont_rep", 1:4, "_filt.bam"),
                            package = pkg)
tss_bamfiles <- system.file("extdata",
                            paste0("cage_rep", 1:3, "_filt.bam"),
                            package = pkg)
pas_bamfiles <- system.file("extdata",
                            paste0("pat_rep", 1:3, "_filt.bam"),
                            package = pkg)
nascent_bamfiles <- system.file("extdata",
                                paste0("planet_rep", 1:2, "_filt.bam"),
                                package = pkg)
```

When running the pipeline on other datasets, use the `list.files(full.names = TRUE)` function instead.

### Usage

The pipeline consists of a few steps which have to be executed sequentially. For a more detailed explanation of the underlying algorithm, see **Appendix 3**.

First of all, attach the libraries which are used at the top level of the example pipeline:

```
library(rtracklayer)
library(magrittr)
```

#### Load BAM files:

The `load_BAM_files()` function takes a vector of BAM filenames as input and imports BAM data into R session as *GRanges* or *GRangesList* objects. The most important argument is `mode`:

- If `mode == "long_reads"`, then the BAM file(s) are expected to contain aligned full-length RNA-seq reads (e.g. ONT Direct RNA-seq reads or HQ consensus reads from PacBio Iso-Seq);
- If `mode %in% c("tss", "pas)`, then aligned 5'- or 3'-tags (sequenced on the Illumina platform in single-end mode) are imported;
- If `mode == "nascent"`, then the BAM file(s) are expected to contain short reads originating from RNA enriched for the nascent fraction. Depending on the method used, the reads may be sequenced in single-end (NET-seq, GRO-seq, TT-seq) or paired-end (pNET-seq, plaNET-seq) mode. In the latter case, the additional `ngs_mode = "PE"` argument has to be specified:

```
long_reads <- load_BAM_files(drs_bamfiles, mode = "long_read")
tss_data <- load_BAM_files(tss_bamfiles, mode = "tss")
pas_data <- load_BAM_files(pas_bamfiles, mode = "pas")
nascent_data <- load_BAM_files(nascent_bamfiles,
                               mode = "nascent", ngs_mode = "PE")
```

Optionally, one may wish to export the loaded NGS data as bedGraph/BED12 tracks for visualization in a genome browser:

```
write_grl_as_bed12(long_reads, "Long_reads.bed")
tss_data %>% merge_GRanges() %>%
  save_GRanges_as_bedGraph("TSS_data_merged.bedgraph.gz")
pas_data %>% merge_GRanges() %>%
  save_GRanges_as_bedGraph("PAS_data_merged.bedgraph.gz")
save_GRanges_as_bedGraph(nascent_data,
                         "Nascent_data_merged.bedgraph.gz")
```

The bedGraph files produced by the `save_GRanges_as_bedGraph()` function are stranded, i.e. encode coverage on the forward (Watson) and reverse (Crick) strands as positive and negative values, respectively. It is important to ensure that the strand information of the input data was processed correctly. If the sequencing coverage is observed always on the opposite strand with respect to known genes, then the strand orientation of such *GRanges* or *GRangesList* object is probably wrong and has to be flipped by the `flip_strand_info()` function.

For faster access to the large BED12 file produced by the `write_grl_as_bed12()` function, we recommend to compress and index it (outside of the R session):

```
file="Long_reads.bed"; bgzip $file && tabix -p bed ${file}.gz && rm $file
```

If using IGV browser, consider to convert the large bedGraph files produced by the `save_GRanges_as_bedGraph()` function to the TDF format:

```
nascent_data %>% seqlengths() %>% as_tibble(rownames = "chr") %>%
  write_tsv("chrom_sizes", col_names = FALSE)
```

```
igvtools totdf Nascent_data_merged.bedgraph.gz Nascent_data_merged.tdf chrom_sizes
```

### Call TSS and PAS

The `call_TCs()` function finds genomic coordinates of transcription start sites (TSS) and polyadenylation sites (PAS) from the 5'- and 3'-tag sequencing data, respectively. It returns a *GRanges* object with numeric `score` metadata column which shows the “strength” of TSS or PAS:

```
tss <- call_TCs(tss_data)
pas <- call_TCs(pas_data)
```

We recommend to visualize the coordinates of called TSS and PAS in a genome browser in parallel with the bedGraph files produced from the input 5’- and 3’-tag sequencing data:

```
rtracklayer::export(tss, "TSS.bed", format = "BED")
rtracklayer::export(pas, "PAS.bed", format = "BED")
```

If by any reason you are not satisfied with the results of peak calling, then changing the default values of `call_TCs` arguments is recommended. In particular, if the input data are noisy (i.e. have high non-specific background), then the results may be improved by using higher `min_tpm`, `q_trim` and `min_score` values. Alternatively, any other peak calling software can be used instead of `call_TCs` (e.g. *CAGEr* or *CAGEfightR*), given that the output is converted to *GRanges* with a valid `score` column.

### Extend long reads towards nearby TSS and PAS

The key idea of the pipeline is to validate and correct full-length RNA-seq reads by orthogonal datasets which detect TSS and PAS with higher accuracy. This idea was implemented in the `extend_long_reads_to_TSS_and_PAS()` function:

```
long_reads_2 <- extend_long_reads_to_TSS_and_PAS(long_reads, tss, pas)
```

### Adjust exon borders

Since long reads from Third generation sequencing platforms in general have higher error rates than the short Illumina reads, the alignments may be relatively inaccurate. In particular, borders of exonic subalignments in long reads are often “fuzzy”. To decrease the number of artifactual alternative exons differing by only a few basepairs, we suggest to use the `adjust_exons_of_long_reads()` function:

```
long_reads_3 <- adjust_exons_of_long_reads(long_reads_2)
```

This step is optional and can be omitted.

### Detect alignment errors

Minimap2, the most popular aligner for Third generation sequencing data, is prone to under-split the full-length RNA-seq reads. Two adjacent exons may appear as a single subalignments in certain reads, whereas as individual subalignments in other reads in the same locus. This effect is most probably due to the relatively low cost for a mismatch between query and template in the long read aligners (as a result, the cost for opening an intronic gap may exceed the cost for erroneously extending the previous exon). This specific kind of alignment artifacts can be found by the `detect_alignment_errors()` function:

```
long_reads_4 <- detect_alignment_errors(long_reads_3)
```

If any exonic subalignment was marked as potential alignment error, then the whole long reads is omitted from the transcript calling procedure. If you trust your aligner, then you can skip the call to `detect_alignment_errors()`.

#### Call transcripts and genes

The corrected and validated long reads are used to call the most probable transcript and gene models by the `call_transcripts_and_genes()` function:

```
out <- call_transcripts_and_genes(long_reads_4)
hml_genes <- out[[1]]
hml_tx <- out[[2]]
fusion_genes <- out[[3]]
fusion_tx <- out[[4]]
reads_free <- out[[5]]
```

The transcripts are classified into High Confidence (HC), Medium Confidence (MC) or Low Confidence (LC) sets, depending on the level of mutual support from the orthogonal methods:

- HC transcripts start in TSS and end in PAS (i.e. are supported by all three datasets);
- MC transcripts either start in TSS, or end in PAS (supported by two datasets, but not by the third one);
- LC transcripts lack both TSS and PAS (supported by the full-length RNA-seq dataset only).

Transcripts with sufficiently strong overlap are combined into genes. Transcripts overlapping two or more called genes are considered fusion transcripts. The function returns a *List* which contains the following elements:

1. HC, MC and LC genes;
2. HC, MC and LC transcripts;
3. Fusion genes;
4. Fusion transcripts;
5. Unused reads which do not overlap with any called gene.

#### Add intervals of nascent transcription

Both the full-length RNA-seq and the 5'- and 3'-tag sequencing methods can only detect mature RNA transcripts. Moreover, they often depend on the poly(A) tail. However, eukaryotic genomes are pervasively transcribed, thus producing many non-coding RNA species which are often transient and/or non-polyadenylated. Such ncRNA species may avoid detection by the conventional RNA sequencing methods, however they become visible in nascent RNA sequencing datasets (NET-seq, GRO-seq, TT-seq, chrRNA-seq etc). In this tutorial we will use plaNET-seq ("plant NET-seq") dataset from Kindgren, Ivanov, and Marquardt (2020), as described above.

The first step is to call continuous intervals of nascent transcription from the imported nascent RNA-seq data by the `call_transcribed_intervals()` function:

```
trans <- call_transcribed_intervals(nascent_data)
transcribed <- trans[[1]]
gaps <- trans[[2]]
```

Alternatively, you can use any other dedicated software instead (e.g. *groHMM*) and coerce its output to a *GRanges* object.

The second step is to interleave the intervals of nascent transcription with the called genes using the `process_nascent_intervals()` function:

```
results <- process_nascent_intervals(hml_genes, transcribed,
                                     tss, pas, reads_free, gaps)
hml_genes_v2 <- results[[1]]
hml_genes_v2_RT <- results[[2]]
lncrna <- results[[3]]
```

The returned *GRanges* object are as follows:

1. The `hml_genes_v2` is almost identical to the input `hml_genes`, except that some MC and LC genes could have found their missing TSS and/or PAS by extension of the gene borders along the adjacent intervals of nascent transcription;
2. The *thick* coordinates of genes in `hml_genes_v2_RT` are identical to the *granges* in `hml_genes_v2` described above. However the *granges* in `hml_genes_v2_RT` represent the union of called genes with their readthrough (RT) tails of nascent transcription;
3. The `lncrna` object contains antisense and intergenic nascent transcripts which are considered as transient and/or non-polyadenylated lncRNAs.

#### Export the results

The results of the pipeline can be exported as BED files for visualization in genome browsers:

```
rtracklayer::export(hml_genes_v2, "Called_genes.bed", format = "BED")
rtracklayer::export(hml_genes_v2_RT,
                    "Called_genes_with_RT_tails.bed", format = "BED")
write_grl_as_bed12(hml_tx, "Called_transcripts.bed")
rtracklayer::export(fusion_genes, "Fusion_genes.bed", format = "BED")
write_grl_as_bed12(fusion_tx, "Fusion_transcripts.bed")
rtracklayer::export(lncrna, "Called_lncRNAs.bed", format = "BED")
```

Observe that the thick parts of genes in the “Called_genes_with_RT_tails.bed” track correspond to the mature transcripts (from TSS to PAS), whereas the thin 3’ extensions denote RT tails. This is different from the traditional visualization scheme (where thick parts are coding exonic intervals between start and stop codons, and thin flanks are 5’ and 3’-UTRs).

### Appendix 1. Strandedness of full-length RNA-seq methods

Full-length RNA-seq on the Oxford Nanopore platform comes in two flavors:

- Direct RNA sequencing:
    - ONT Direct RNA Sequencing kit;
- cDNA sequencing:
    - ONT cDNA-PCR Sequencing kit;
    - ONT Direct cDNA Sequencing kit;
    - A third party cDNA synthesis kit (e.g. Takara SMARTer PCR cDNA synthesis kit) + ONT Ligation Sequencing kit.

In the case of Direct RNA-seq, ONT sequencing adapters are ligated only to the 3'-end of the original RNA molecule. Sequencing always occurs in the 3' -> 5' direction. Therefore, the long reads retain the strand orientation of the original RNA molecules.

In the case of full-length cDNA sequencing, ONT sequencing adapters are ligated to both ends of the double-stranded cDNA. It is impossible to predict direction of the sequencing (the first or the second strand of cDNA can be sequenced with equal probabilities). In other words, the strand orientation of the original RNA molecule is lost. The *TranscriptomeReconstructoR* assumes that the input long reads are properly stranded, thus it cannot be used with unstranded cDNA sequencing data.

Theoretically, it might be possible to recover the strand orientation of a full-length cDNA dataset, given that the cDNA synthesis method produces asymmetric cDNA flanks (i.e. the oligo-dT primer and the template switching oligo have different 5' sequences). Unfortunately, it is not the case of the commonly used Takara (Clontech) SMARTer cDNA synthesis kit (where both the 3' SMART CDS Primer II A and the SMARTer II A Oligonucleotide begin with the same 5'-AAGCAGTGGTATCAACGCAGAGTAC-3' sequence). However, if a custom oligo-dT primer were used instead of the 3' SMART CDS Primer II A, then the strandedness of long reads could be guessed from the adapter sequences by a custom script.

Luckily, the PacBio Iso-Seq pipeline automatically detects the strand orientation of HQ consensus reads (even despite the fact they are obtained from double-stranded cDNA).

In conclusion, long reads from ONT Direct RNA-seq and PacBio Iso-Seq are stranded and thus can be used as input for the *TranscriptomeReconstructoR*. Long reads from ONT cDNA-seq are unstranded and thus are not compatible with *TranscriptomeReconstructoR* (unless the strand orientation was recovered by a custom approach).

### Appendix 2. Annotation-guided mode

The pipeline described above was designed for *de novo* calling of gene and transcript models from the experimental data only. However, in some cases an existing annotation may help to refine the called models. For example, MC and LC transcripts lack TSS and/or PAS. One possible reason is that all long reads in given locus were obtained from RNA fragments, not from full-length RNA molecules. Thus, the distance between the borders of long reads and the relevant TSS or PAS exceeds the reasonable limits (which are controlled by `read_flanks_up` and `read_flanks_down` arguments of the `extend_long_reads_to_TSS_and_PAS()` function). In this case, the existing annotation may help to connect the TSS/PAS with the called gene. The truncated called transcript will be extended along its best mate among the annotated transcripts towards the called TSS and/or PAS.

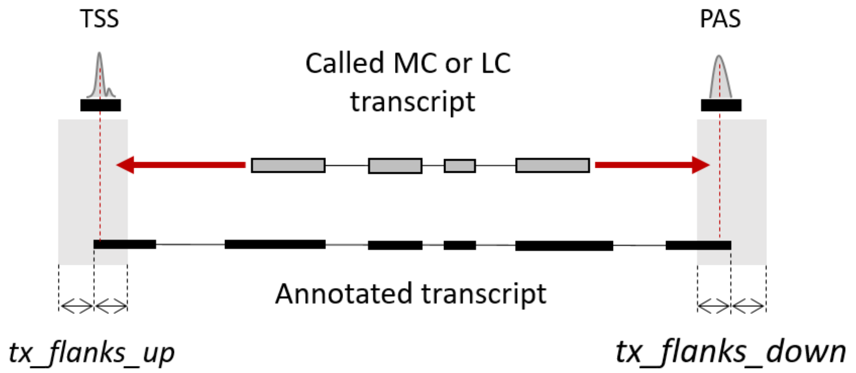


Another possible scenario is that a low expressed gene has both TSS and PAS, however zero coverage in the full-length RNA-seq dataset (due to relatively low throughput of Third generation sequencing). If these TSS and PAS are connected by an annotated gene, then such gene and its transcripts can be imported into the called gene and transcript models.

Finally, the data-driven correction of exon borders by the `adjust_exons_of_long_reads()` function does not guarantee that the coordinates of splice sites were detected precisely (the majority vote does not always provide the right decision). On the other hand, the existing annotations are usually based on short read RNA-seq data which are characterized by low error rate and thus high precision split alignments. Therefore, the *de novo* detected exon borders can be further corrected by the annotated splice sites.

The `refine_transcripts_by_annotation()` function implements these three ideas. It should be used after calling *de novo* models by the `call_transcripts_and_genes()` function, but **before** running the `process_nascent_intervals()` function. It needs the following input objects:

- HC, MC and LC transcripts (*GRangesList* returned by the `call_transcripts_and_genes()` function);
- Annotated exons grouped by transcript (*GRangesList* object, e.g. returned by the `exonsBy(by = "tx")` function);
- TSS and PAS (*GRanges* objects returned by the `call_TCs()` function);
- Fusion transcripts (*GRangesList* object returned by the `call_transcripts_and_genes()` function).

```
library(TxDb.Athaliana.BioMart.plantsmart28)
txdb <- TxDb.Athaliana.BioMart.plantsmart28
annot_exons <- exonsBy(txdb, by = "tx")

ref <- refine_transcripts_by_annotation(hml_tx, annot_exons,
                                        tss, pas, fusion_tx)
hml_genes <- ref[[1]]
hml_tx <- ref[[2]]
fusion_genes <- ref[[3]]
fusion_tx <- ref[[4]]
```

The `refine_transcripts_by_annotation()` function returns a *List* with the following elements:

1. Updated HC, MC and LC genes (*GRanges* object);
2. Updated HC, MC and LC transcripts (*GRangesList* object);
3. Updated fusion genes (*GRanges* object);
4. Updated fusion transcripts (*GRangesList* object).

If nascent RNA-seq dataset is used in the study, then the updated HC, MC and LC genes are used as input for the `process_nascent_intervals()` function. If not, then the objects returned by the `refine_transcripts_by_annotation()` function are the final gene and transcript model to be exported into BED files.

### Appendix 3. The algorithm

#### call_TCs()

The input for this function is *GRangesList* object, where each element is *GRanges* with stranded 5'- or 3'-tag counts in a biological replicate sample. By default, only genomic positions with non-zero tag count in all replicate samples are considered. This condition can be relaxed by setting `min_support` argument to a value smaller than the number of input BAM files. After filtering by minimal support, the replicates are pooled, the signal is normalized to tags per million (TPM), and the tag clusters (TCs) are called as continuous intervals with the signal above the `min_tpm` threshold. Adjacent TCs are merged together, if the distance between them does not exceed `max_gap` bp.

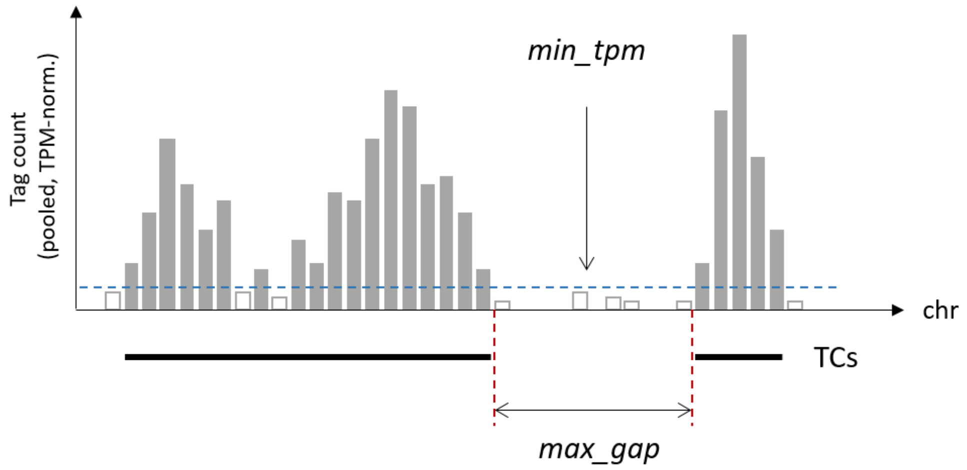


Then TCs are quantile trimmed to skip the trailing tails of weak signal which often surround the TC summit. The amount of trimming is controlled by the `q_trim` argument.

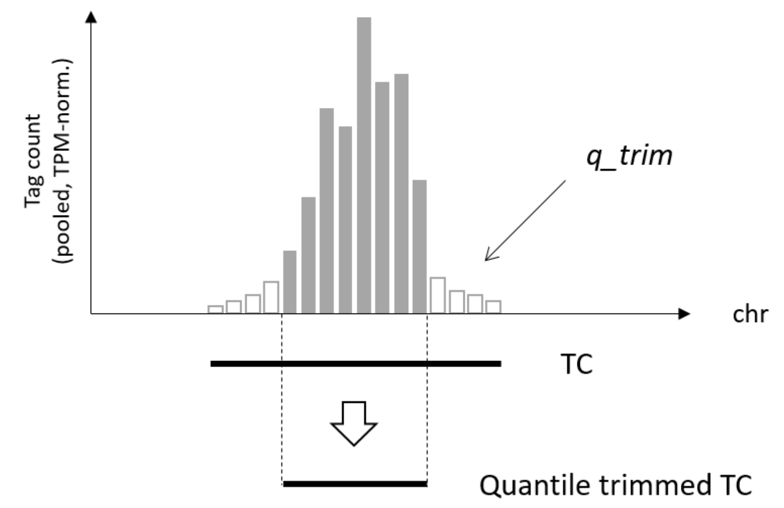


After trimming, the TCs are merged again (with the same `max_gap` parameter). Finally, each TC gets a score which is the TPM-normalized signal averaged between the replicates. TCs with scores less than `min_score` are skipped. The remaining TCs are returned as *GRanges* object with `score` metadata column.

#### extend_long_reads_to_TSS_and_PAS()

This function takes *GRangesList* with long reads as input. For each long read, two genomic windows are generated:

- Upstream window: `read_flanks_up` bp around 5’ end of the long read;
- Downstream window: `read_flanks_down` bp around 3’ end of the long read.

The functions searches for called TSS overlapping with the upstream window, and for called PAS overlapping with the downstream window. If multiple TSS or PAS were found, then the “strongest” of them (the one with the highest score) is chosen. Then the long reads are extended towards the summits of the found TSS and PAS.

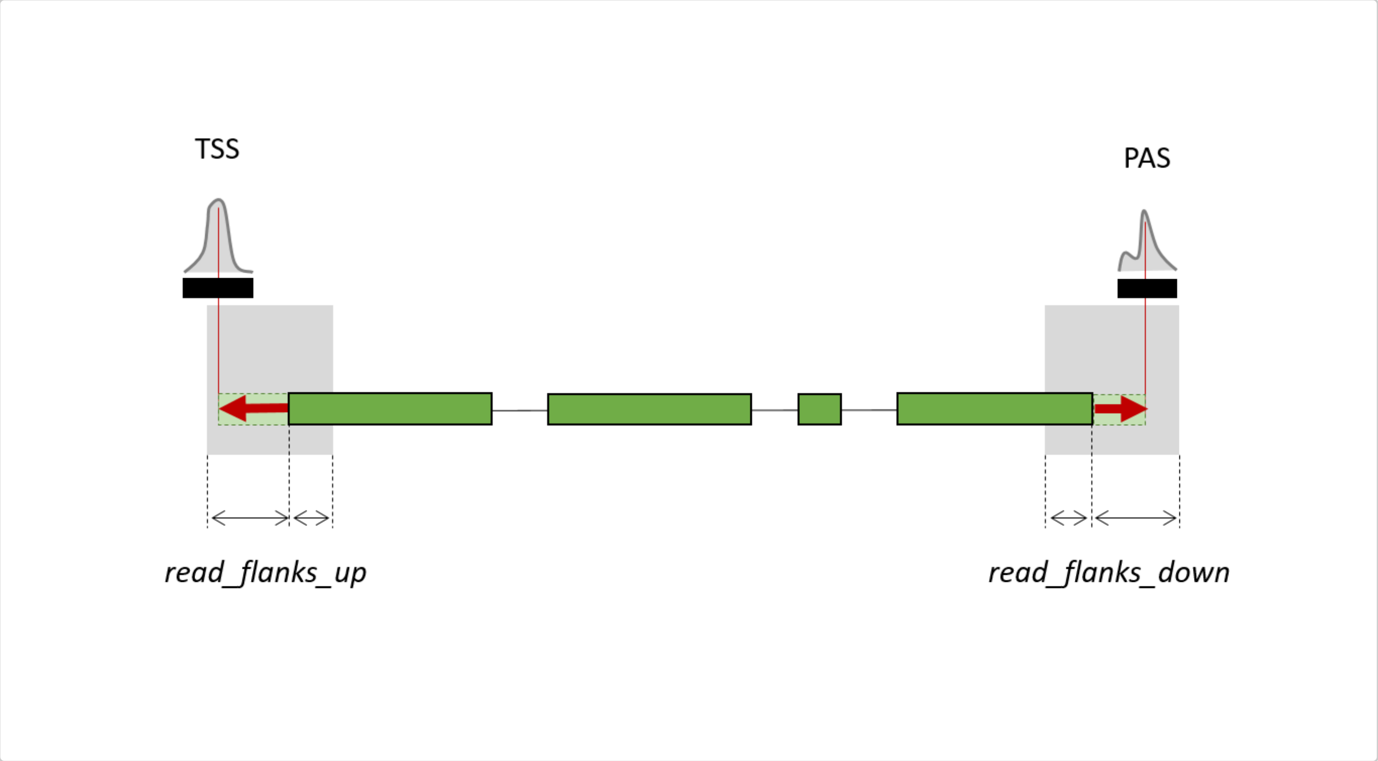


Extended long reads which start in a TSS and end in a PAS are classified as “complete”, otherwise as “truncated”. If a locus (defined as an overlapping group of reads) does not contain any complete read, then the longest truncated read is re-classified as “complete”, and its free end is considered as the best estimate of the transcript border. In addition, other truncated reads in this locus which can be safely extended along the “guide exon”, are also re-classified as “complete”.

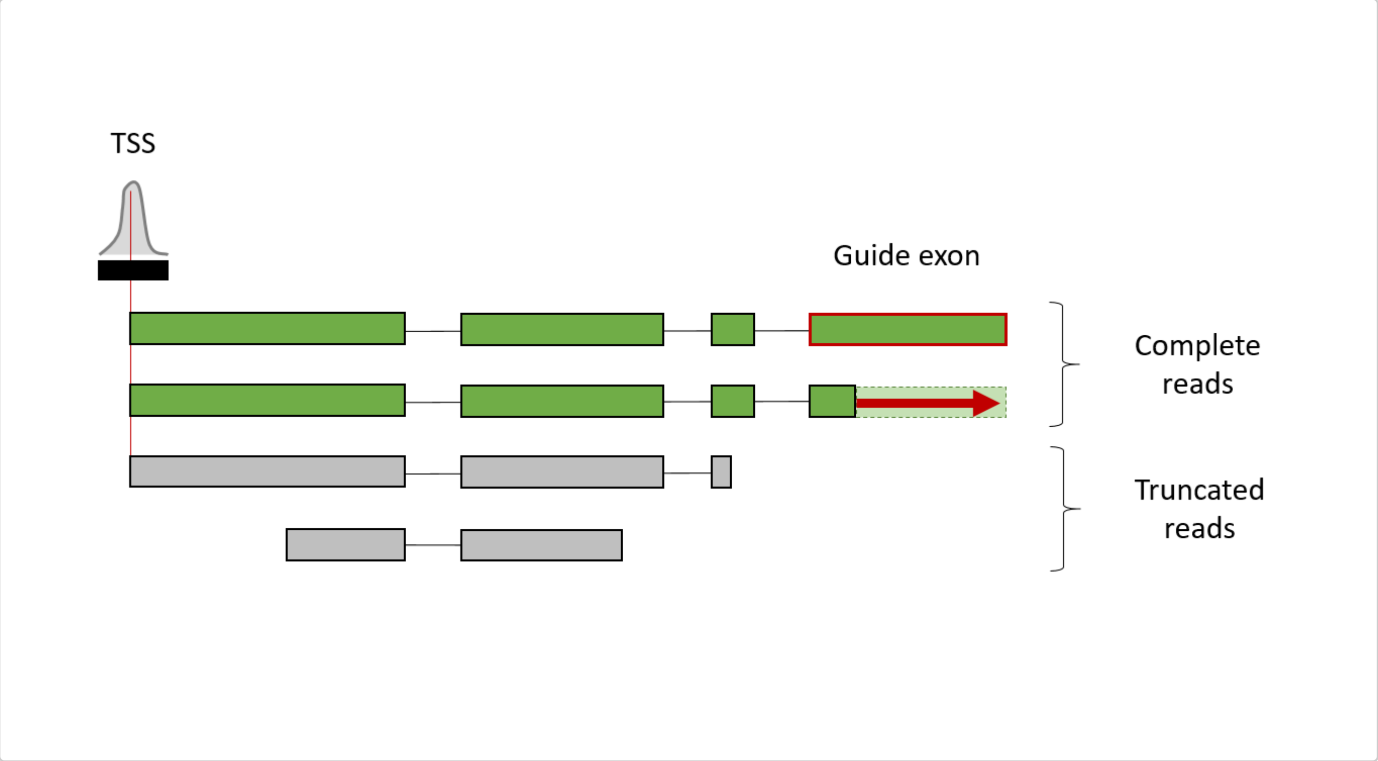


The function returns *GRangesList* object containing extended long reads with additional metadata columns. Only reads marked as “complete” are used for the transcript calling procedure (see below).

#### adjust_exons_of_long_reads()

This function extracts all splice sites from the input long reads, clusters them within `max_exon_diff` bp distance and then unifies the borders of exonic subalignments to the most frequently observed coordinate within each cluster.

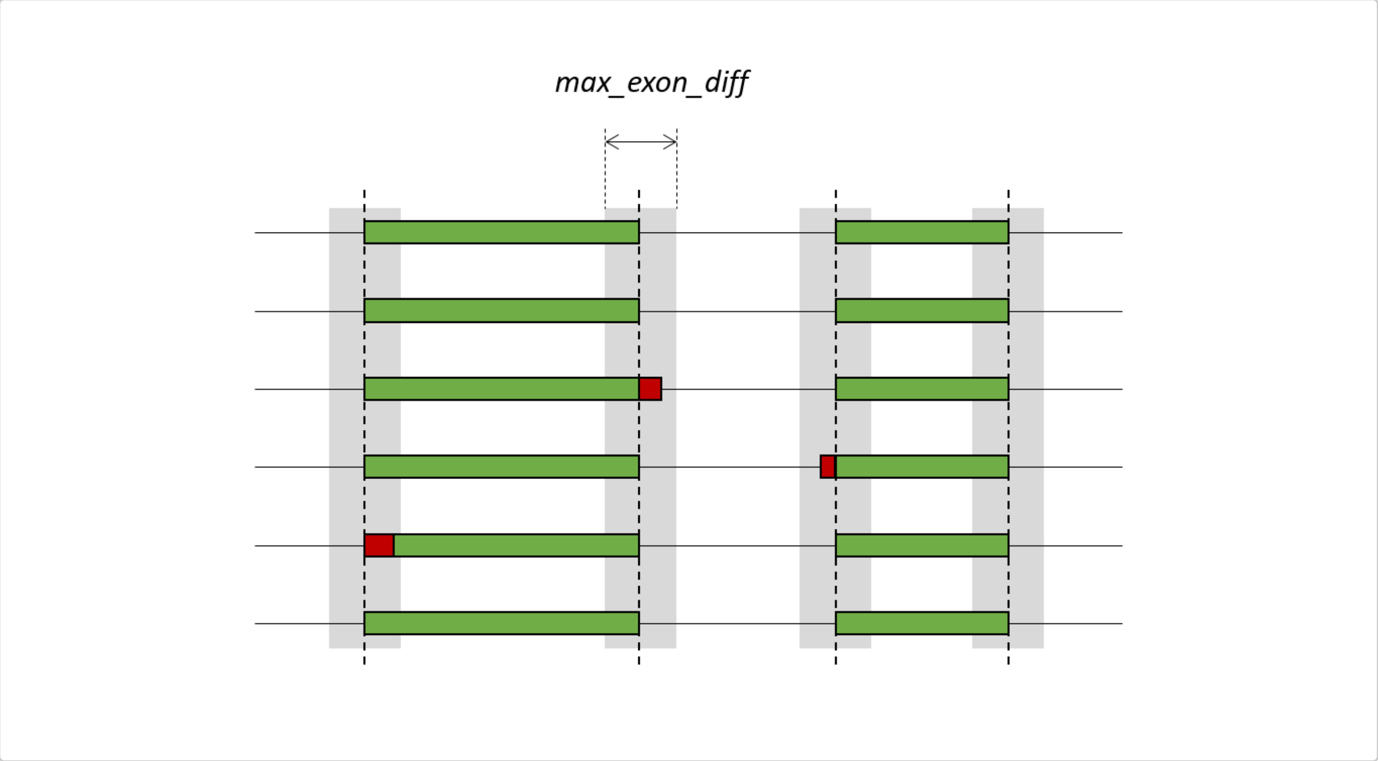


Both input and output for this function are *GRangesList* objects containing long reads with identical metadata columns but different coordinates of subalignments. Observe that this “majority vote” procedure does not guarantee to find the true splice sites. However, the number of artifactual transcript isoforms with alternative 5’- or 3’-splice sites differing by only a few basepairs is expected to decrease significantly.

#### detect_alignment_errors()

Probably the most common type of alignment errors in full-length RNA-seq reads in under-splitting, i.e. erroneous extension of an exon over the adjacent intronic region. As a result, the next exon appears as missing from such read (although present in other reads aligned within the same locus). Assuming that the majority of reads still align correctly, we suggest to detect such alignment errors by comparing each subalignment to the set of constitutive exons.

The constitutive exons are defined here as exonic intervals supported by more than 50% of reads in given locus (but not less than reads). Subalignment with an alternative 5’- and/or 3-border (relative to a constitutive exon) are considered valid alternative exons, only if the next constitutive exon is also present in the read. Otherwise, the alignment is marked as a possible alignment error.

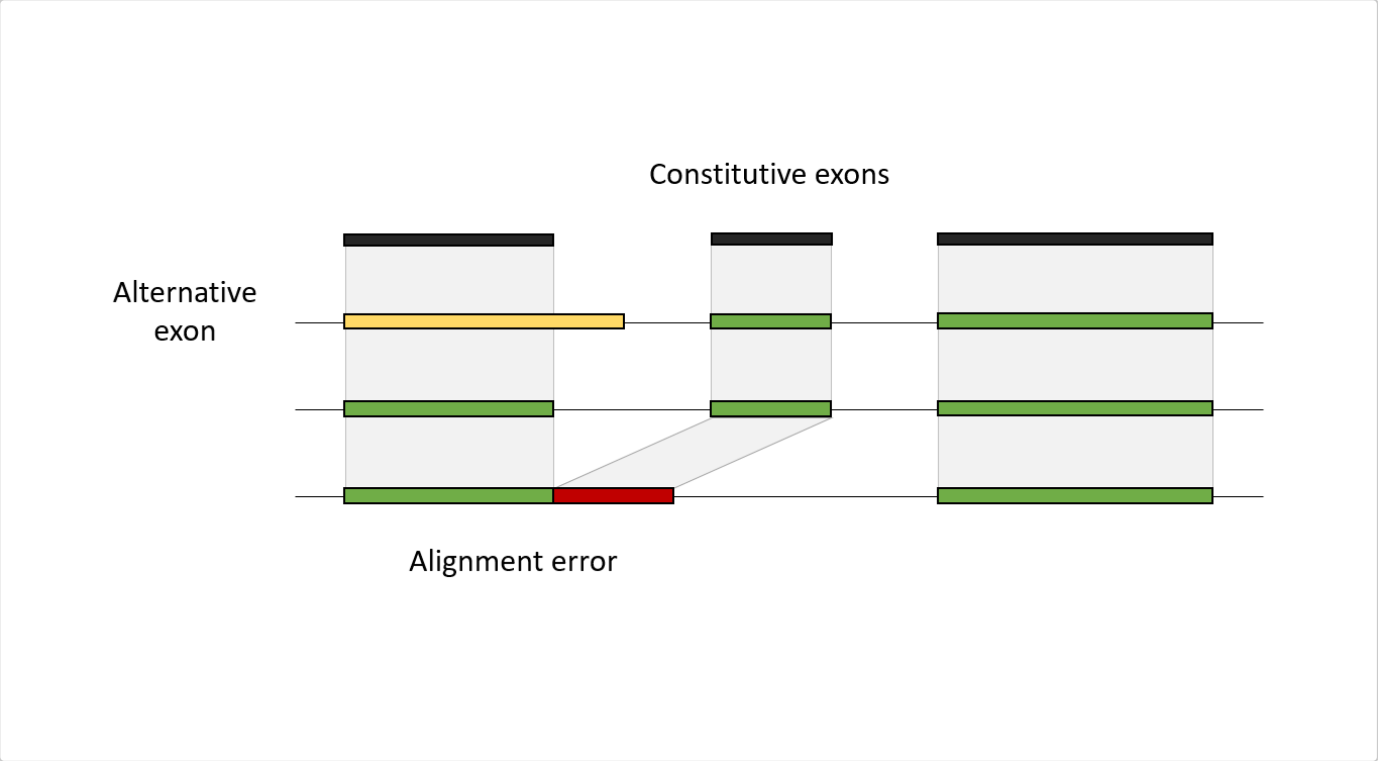


Both input and output for this function are *GRangesList* objects containing long reads with identical coordinates of subalignments. The output object has a new metadata column which determines if the subalignment is a suspected alignment error.

#### call_transcripts_and_genes()

The input for `call_transcripts_and_genes()` function is *GRangesList* object containing long reads. The long reads are expected to be already processed by the upstream functions in the pipeline (`extend_long_reads_to_TSS_and_PAS()`, `adjust_exons_of_long_reads()` and `detect_alignment_errors()`). These upstream steps serve two purposes: i) attenuate the alignment “noise” associated with Third generation sequencing data; ii) detect reads which are not informative of the original transcripts (i.e. reads truncated at one or both ends and/or containing suspected alignment errors). Such reads are skipped from consideration when calling the transcripts and genes. Only “complete” reads without misaligned exons are used. Observe that some reads marked as “complete” may still miss TSS and/or PAS (see the description of `extend_long_reads_to_TSS_and_PAS()` function above).

Transcript calling is a multi-step procedure:

1. At the first step, we use only reads which start in a TSS and end in a PAS. Identical reads are collapsed into isoforms of High Confidence (HC) transcripts. An HC transcript is supported by all three methods (full-length RNA-seq + 5’ tag sequencing + 3’ tag sequencing) and therefore has reliable outer borders;
2. Strongly overlapping HC transcripts (intersection/union >= `clust_threshold`) are combined into HC genes;
3. Transcripts which overlap two or more adjacent genes by at least `min_overlap_fusion` fraction if their lengths, are considered fusion transcripts and separated from the transcript pool. Fusion transcripts result from inefficient transcription termination at PAS sites;
4. Find “free” reads which satisfy the following conditions:
    - Were not used for calling HC transcripts;
    - Do not overlap with any called HC transcript by more than `max_overlap_called` fraction of either transcript length or read length;
    - Are not shorter than `min_read_width` bp;

5. Among the “free” reads, find those which either start in a TSS, or end in a PAS (but not both). There reads are used to call Medium Confidence (MC) transcripts. An MC transcript is supported by only two methods (full-length RNA-seq + either 5’- or 3’ tag sequencing). Thus, only one outer border of an MC transcript is reliable, whereas the other may be truncated;
6. Repeat steps 2-4 (gene calling, fusion transcript detection, updating “free” reads) for the MC transcripts;
7. Similarly, call Low Confidence (LC) transcripts from “free” reads which lack both TSS and PAS. An LC transcript is supported by full-length RNA-seq only and thus may be truncated from both ends;
8. Within each HC, MC or LC gene, we skip the minor isoforms (which collectively represent not more than `skip_minor_tx` fraction of all supporting reads in this locus);
9. Finally, the HC, MC, LC and fusion genes are recalculated and enumerated.

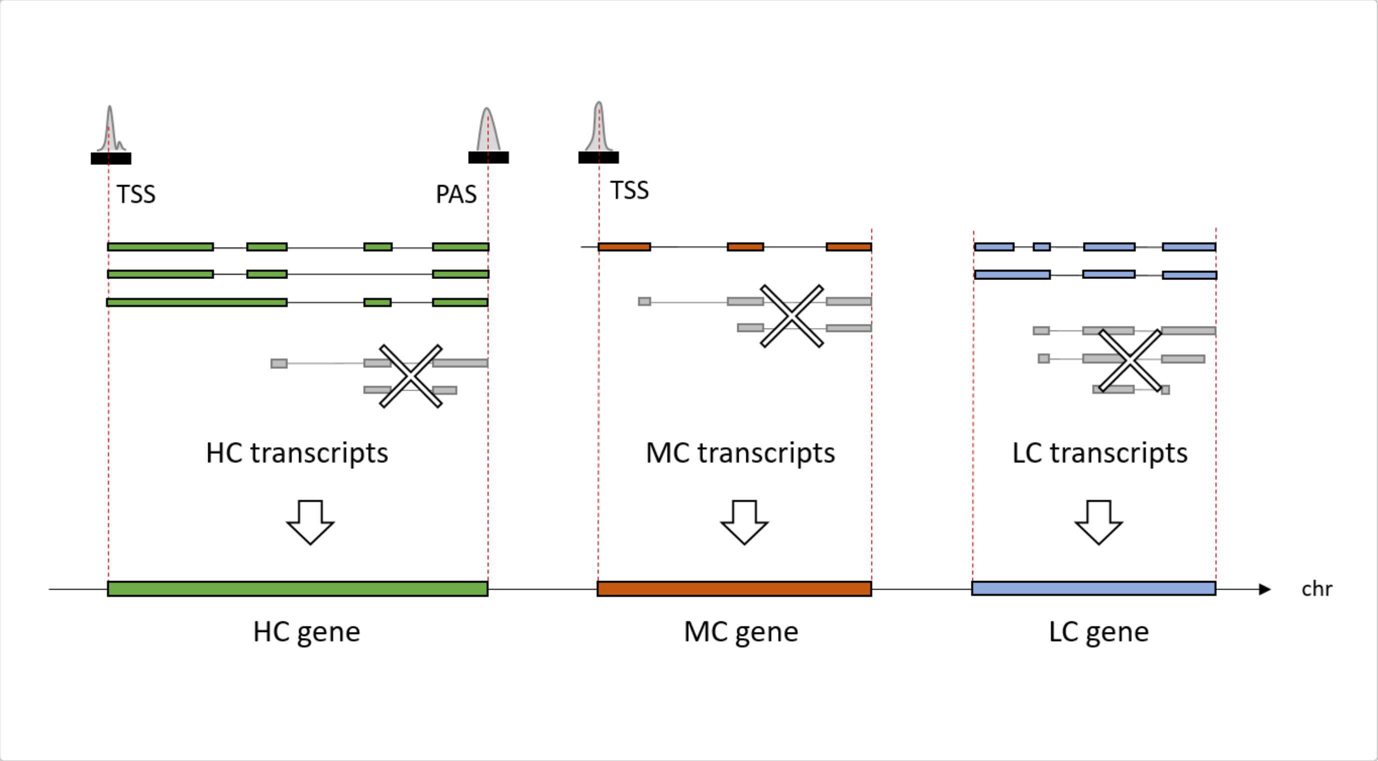


The MC and LC transcripts are called because some loci may have full-length RNA-seq reads, however lack sufficiently strong TSS and/or PAS tag clusters in close proximity to outer borders of the long reads. The mismatch between different methods can happen due to various reasons, e.g.:

- Library preparation, sequencing or alignment biases of 5’- or 3’-tag sequencing methods which result in low sensitivity of TSS or PAS detection in given locus;
- Fragmentation of the original RNA molecules (either *in vivo* or during the full-length RNA-seq library preparation) which results in the excess of truncated long reads;
- Using datasets obtained from different biological samples.

The latter possibility can be easily avoided by properly planning the experimental workflow (one should always prepare libraries for the different sequencing methods from the same biological sample). However, the former two sources of unwanted variability cannot be fully excluded. Thus, the MC and LC transcripts, although having potentially inaccurate outer borders, serve as the best estimates for the transcriptional activity in such loci. The iterative transcript calling procedure ensures that the MC transcripts can be called only outside of the HC loci, and LC transcripts can be called only outside of both HC and MC loci.

Long reads either marked as truncated by , or containing a misaligned exon (as revealed by ), are skipped from the transcript calling procedure. The remaining long reads are collapsed into transcripts. The transcripts are classified into high confidence (HC), medium confidence (MC) and low confidence (LC) groups: This iterative procedure of transcript calling ensures that highly expressed HC loci are not contaminated with less reliable MC or LC transcripts. The MC and LC transcripts can be called only in those loci where the full-length reads are lacking. To decrease the risk of picking up products of partial RNA degradation, MC and LC transcripts can be called only from reads longer than bp. The called HC, MC and LC transcripts are clustered into HC, MC and LC genes, respectively. A pair of transcripts of the same type having overlap (intersect/union) above the are considered belonging to the same gene. Within each gene, the minor transcripts (collectively representing up to fraction of the supporting reads) are skipped. This is done to decrease the risk of calling artifact alternative transcripts in highly expressed genes (assuming that undetected alignment errors are relatively rare events). This behavior can be suppressed by setting . Finally, transcripts which overlap at least two other disjoint transcripts by at least fraction of their lengths, are considered fusion transcripts.

The `call_transcripts_and_genes()` function returns a *List* of *GRanges* and *GRangesList* objects which contain the called HC, MC and LC genes and transcripts, fusion genes and transcripts, as well as the “free” reads.

### call_transcribed_intervals()

This function takes input *GRanges* object with `score` metadata column which represents the stranded coverage in a nascent RNA sequencing experiment. The function simply finds intervals with sequencing coverage not less than `min_signal`. Adjacent intervals (separated by low coverage gaps not exceeding `max_gapwidth` bp) are merged. Intervals shorter than `min_width` are skipped.

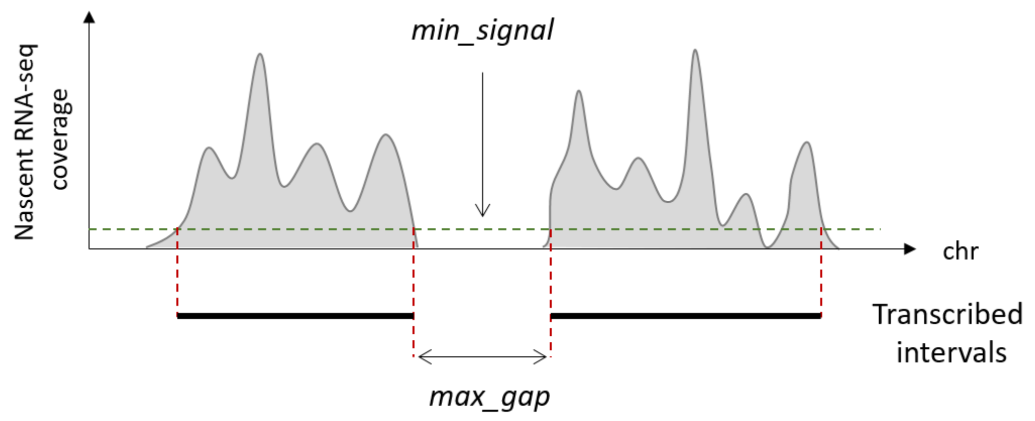


The return value is a *List* of two *GRanges* objects which contain the continuous transcribed intervals and the low coverage gaps, respectively.

### process_nascent_intervals()

This function takes a few inputs *GenomicRanges* objects:

- HC, MC and LC genes (*GRanges* returned by the `call_transcripts_and_genes()` function);
- Intervals of nascent transcription (*GRanges* returned by the `call_transcribed_intervals()` function);
- TSS and PAS (*GRanges* returned by the `call_TCs` function);
- (optionally) Unused reads (*GRangesList* returned by the `call_transcripts_and_genes()`);
- (optionally) Low coverage gaps (*GRanges* returned by the `call_transcribed_intervals()`).

All called genes overlap with intervals of nascent transcription, at that the transcribed intervals are usually wider than the genes. In particular, many genes have trailing “tails” of nascent transcription (known as readthrough, or RT, tails) immediately downstream from their PAS. The `process_nascent_intervals()` function combines the called genes with their RT tails. The outer borders of HC, MC and LC genes are extended to cover the whole transcription unit (at that, the original gene coordinates which correspond to the area of productive elongation are moved to the *thick* metadata column of the *GRanges* object).

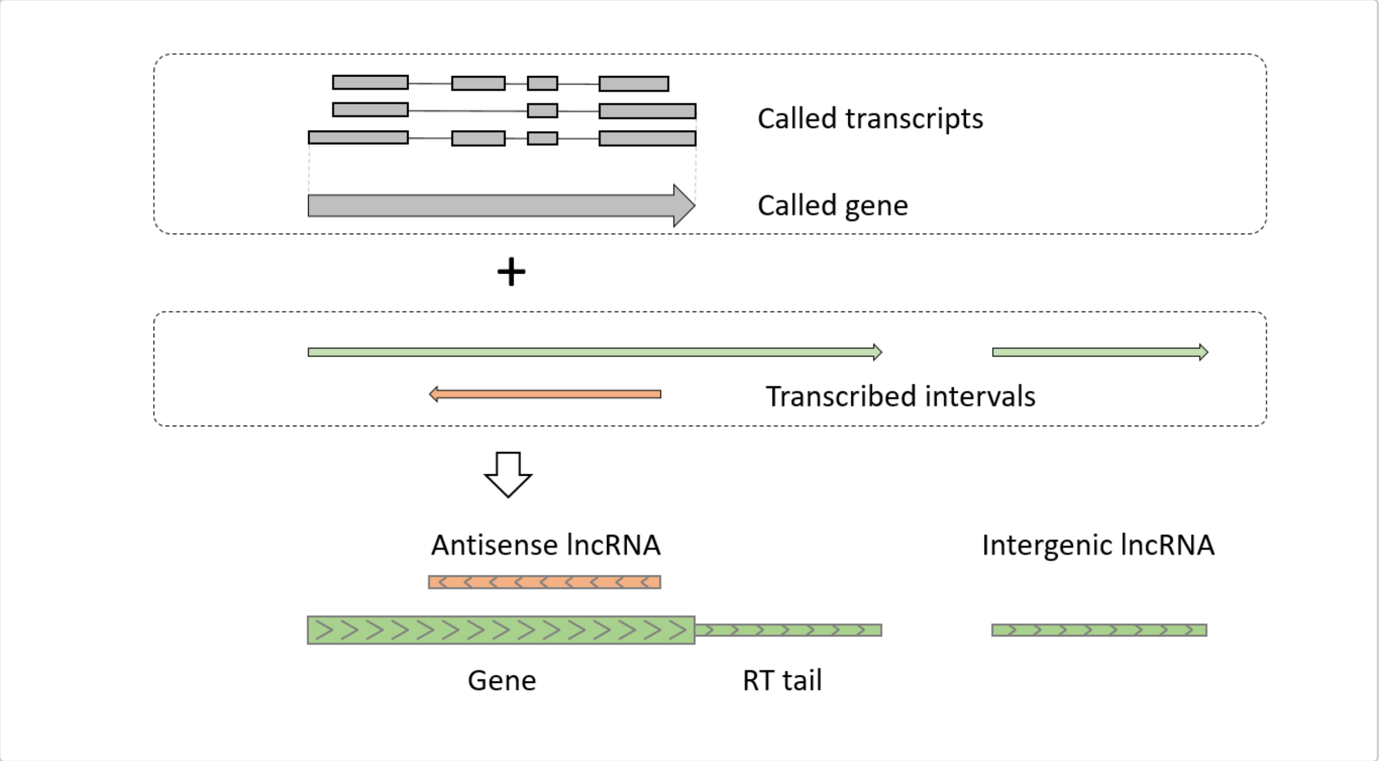


In addition, the area of productive elongation in MC and LC genes can be expanded along the up- and/or downstream intervals of nascent transcription, if they connect the gene to a sufficiently strong TSS and/or PAS (this behavior is controlled by the `extend_along_nascent`, the `extension_flanks` and the `min_score_2` arguments).

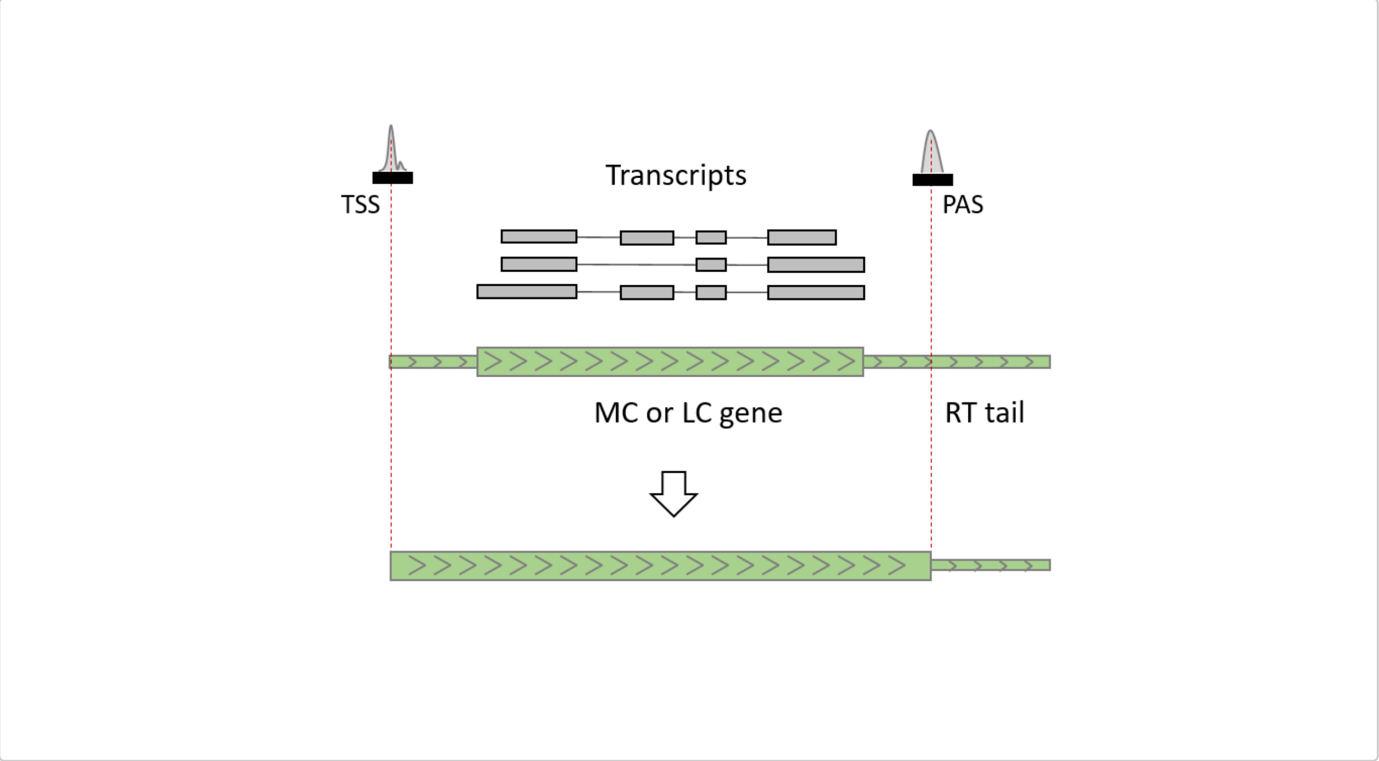

Finally, intervals of nascent transcription which do not overlap with any called gene on the same strand and are longer than `min_lncrna_width`, are considered long non-coding RNAs (lncRNAs). If they overlap with strong TSS and/or PAS (the minimal TC score is controlled by the `min_score_2` argument), they are further refined (extended or split) to match the transcription initiation and termination landscape.

The `process_nascent_intervals()` function returns a *List* with the following elements:

1. HC, MC and LC genes (*GRanges* almost identical to the input `hml_genes` object, except that some genes might have found their missing TSS and/or PAS);
2. HC, MC and LC genes decorated with the RT tails (*GRanges* object with the *thick* metadata column);
3. Antisense and intergenic lncRNAs (*GRanges* object).
