## Supplementary figures and images for "TrancriptomeReconstructoR: data-driven annotation of complex transcriptomes"

### Supplementary Figures S1-S4

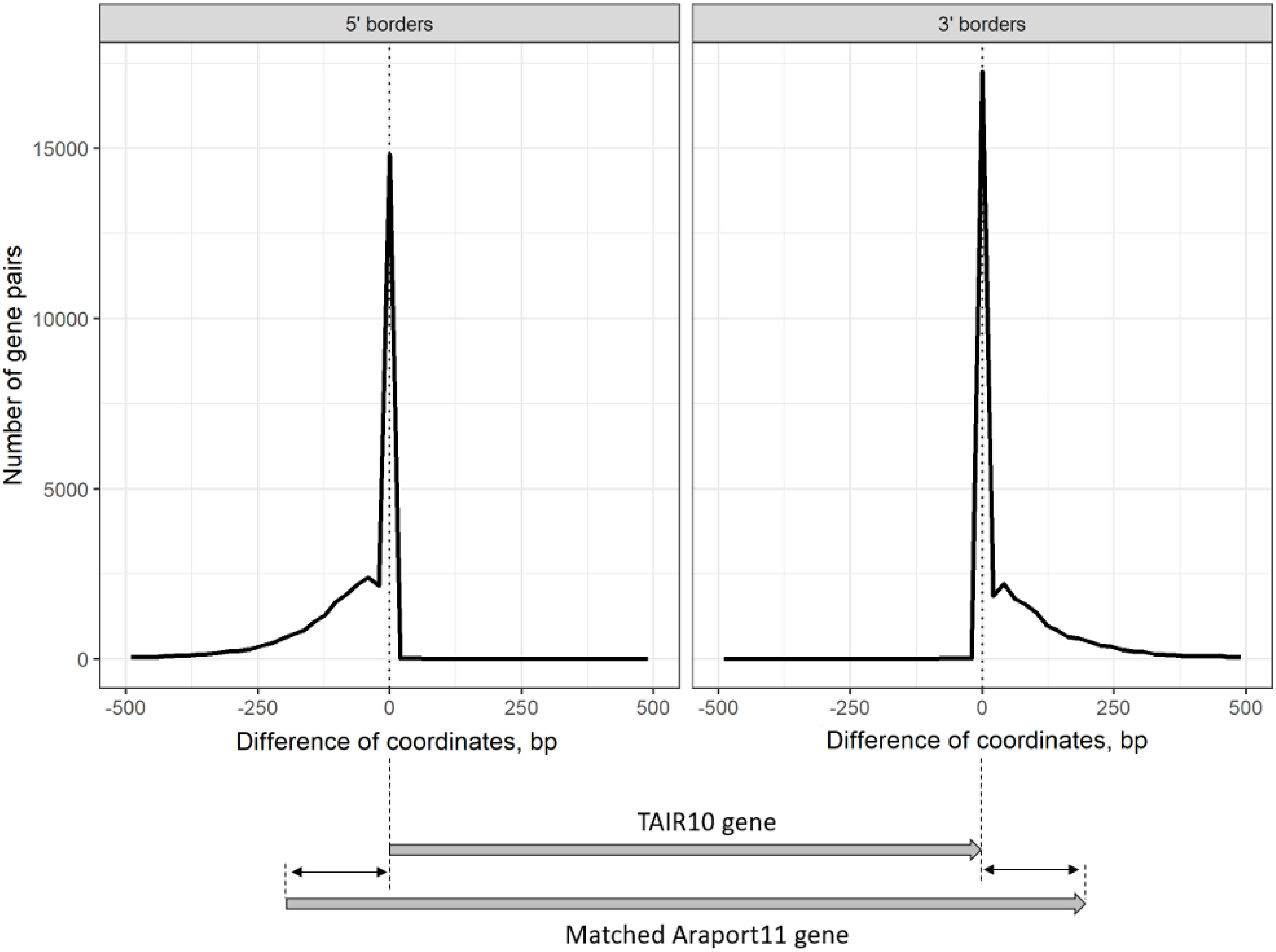

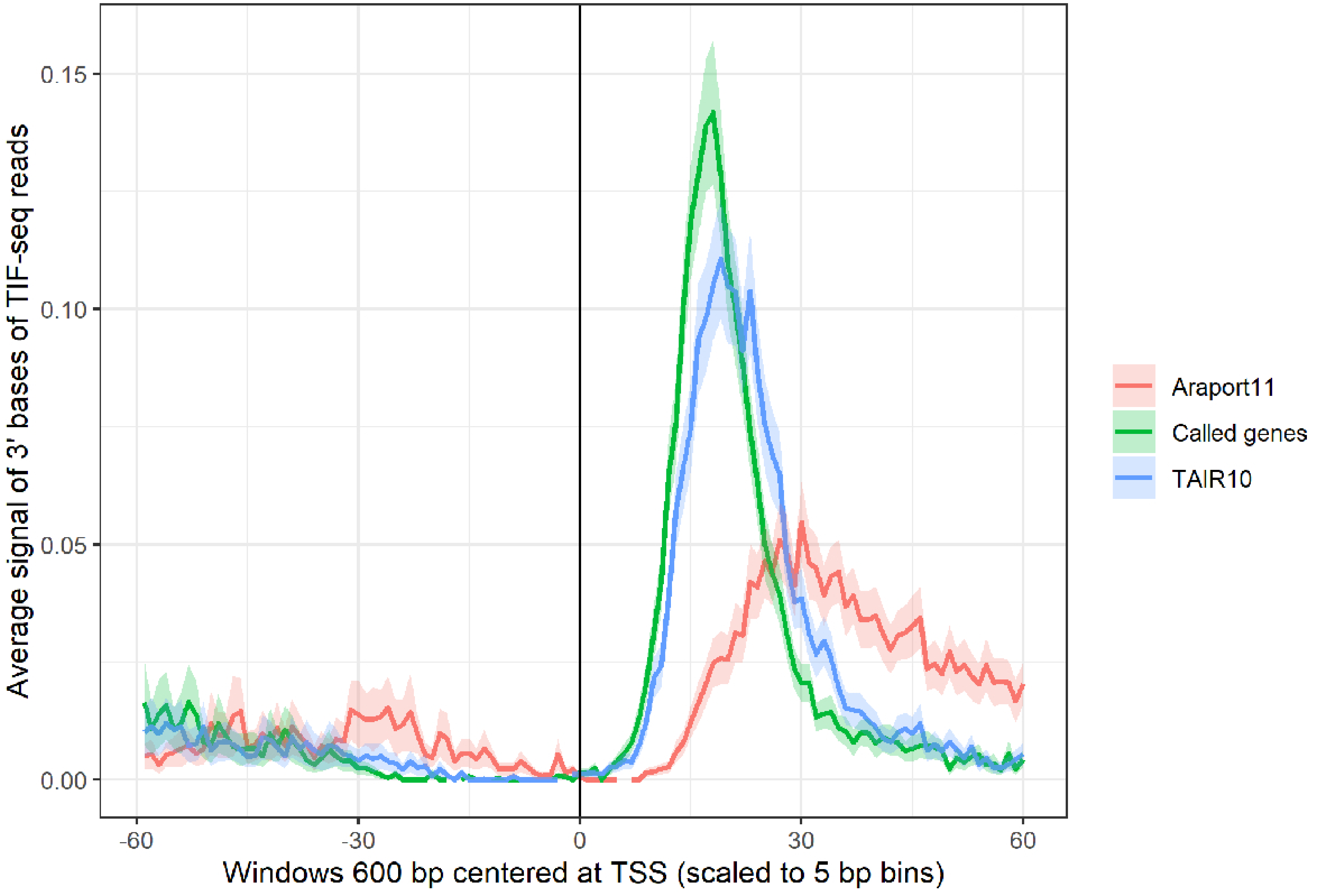

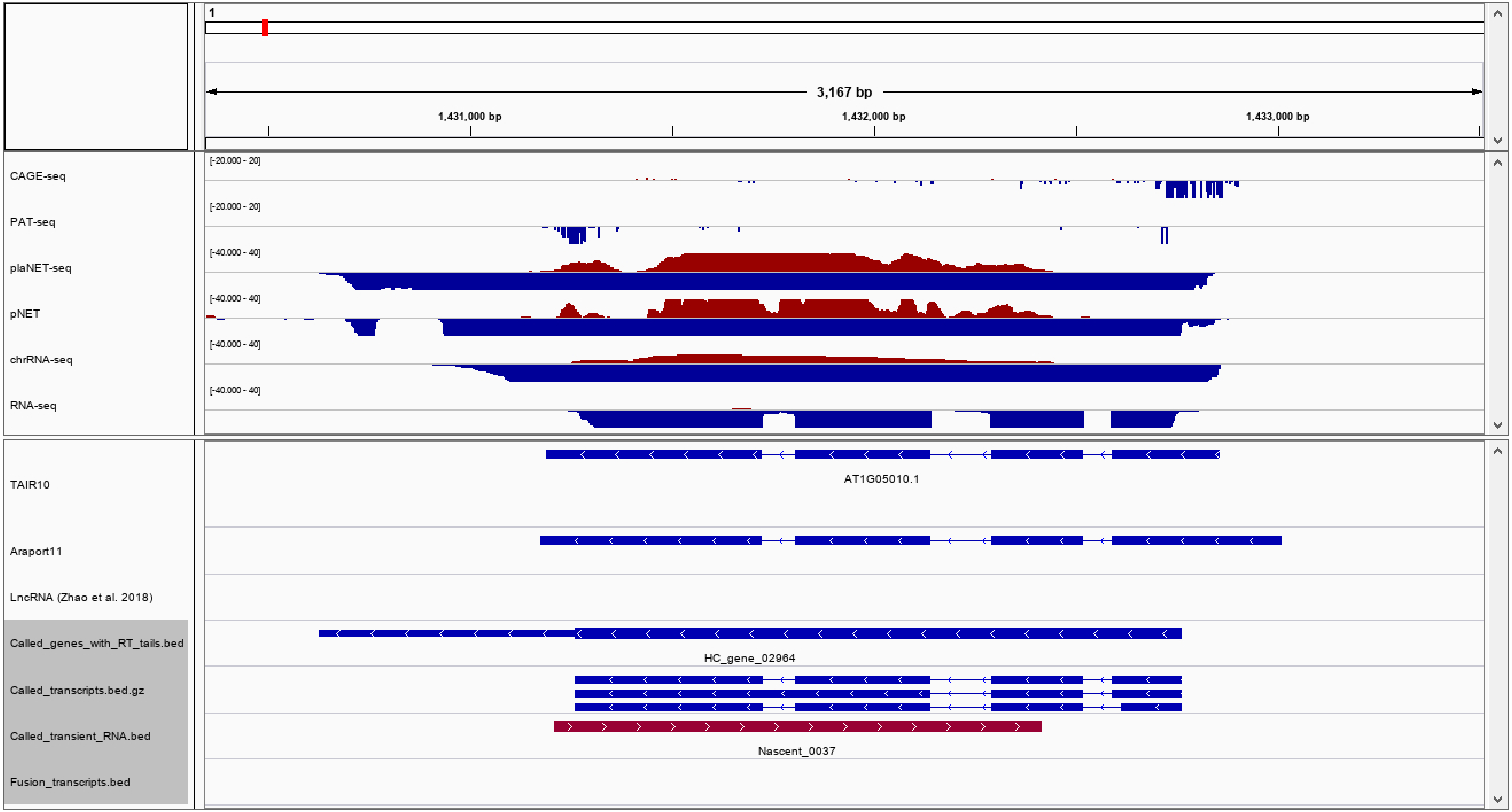

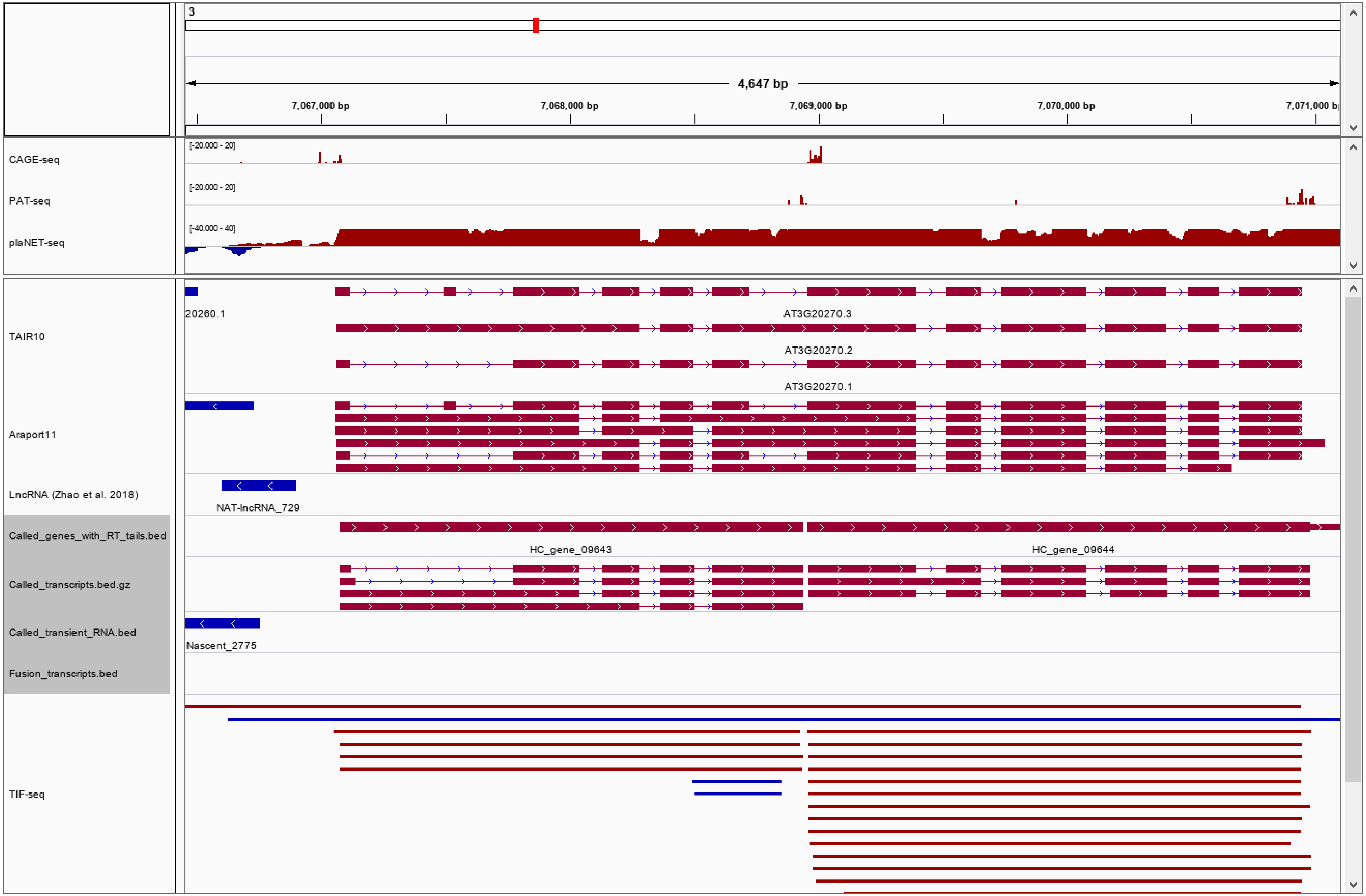
